# A Cross-Study Multi-Organ Cell Atlas of *Macaca fascicularis* Informed by Human Foundation Model Annotation: A Resource for Translational Target Assessment

**DOI:** 10.64898/2026.03.17.711997

**Authors:** Terezinha M. Souza, Joshua T. Gamse, Leandro Moreno, Marilijn van Rumpt, Gonzalo Nuñez-Moreno, Indu Khatri, Saskia D. van Asten, Nichal V. Khusial, Empar Baltasar-Pérez, Ragini Adhav, Tamim Abdelaal, Anna Wojtuszkiewicz, Jorg J. A. Calis, Attila Csala, Anna Dahlman, Claudette L. Fuller, Craig J. Thalhauser, Iris C.R.M. Kolder

## Abstract

Non-human primates (NHPs), particularly *Macaca fascicularis* (cynomolgus macaque), represent an essential model for preclinical assessment of biologics due to their high genetic and physiological similarity to humans. However, mounting regulatory pressure to reduce NHP use and the lack of a unified, well-annotated single-cell atlas currently limits both target qualification and mechanistic interpretation of toxicity in this species. To address this gap, we assembled and harmonized the largest single-cell transcriptomic atlas of *M. fascicularis* to date, integrating 30 publicly available studies spanning 57 anatomical regions, 43 organs and 14 physiological systems. We implemented a scalable framework for cross-species cell type annotation by embedding both cynomolgus monkeys and human (Tabula Sapiens V2) datasets into a shared reference space using Universal Cell Embeddings (UCE), enabling consistent harmonization of cell identities. In total, 27 organs were annotated using human reference labels, while the remaining sets retained author-provided annotations or labels transferred from other cynomolgus studies with available annotations. The resulting atlas comprises over 2.5 million cells and demonstrates concordance in cell-type-specific expression patterns between cynomolgus and humans, including tissue-specific markers and targets relevant for biologics development. Through translational use cases, we illustrate how this resource can be applied to assess target expression in tissues affected by concordant human-NHP toxicities, investigate ocular adverse events associated with antibody-drug conjugates (ADCs), and identify species-specific features of immune cell subtypes with known safety implications. By enabling scalable, high-resolution, cross-species comparisons of gene expression across organs, tissues, and cell states, this atlas supports improved target qualification, more mechanistic interpretation of toxicities, and evidence-based decisions on the relevance and design of NHP studies. Collectively, this work provides a unified cross-species single-cell resource for cynomolgus monkey and a modular computational framework that advances new approach methodologies and contributes to the refinement and reduction of NHP use in preclinical research.

## Introduction

In preclinical drug development, the non-human primate (NHP) model *Macaca fascicularis* (also known as crab-eating macaque, cynomolgus macaque or cyno) is regarded as an important model to assess the safety of biologics (Iwasaki et al. 2019; Ménochet et al. 2022). Given its high genetic similarity to humans, the cynomolgus model exhibits developmental, physiological and pathological responses comparable to humans (Qu et al. 2022) and among commonly-used preclinical models, it has demonstrated the most frequent cross-reactivity to monoclonal antibodies (mAbs) (Iwasaki et al. 2019). Over the past decades, the use of target biologics has expanded and diverged into a range of modalities, including monoclonal antibodies, bispecifics, Fc-fusion proteins and antibody-drug conjugates (ADCs), which in turn has increased the demand for safety studies in the cynomolgus model.

In recent decades, concerns over the use of animals in research have grown, leading to health authority and international guidance to reduce the use of animals in preclinical research. The 3Rs (replacement, refinement, and reduction) initiatives and the UK’s National Centre for the Replacement, Refinement and Reduction of Animals in Research (NC3Rs) have been proactive advocates for rigor in animal welfare and use for more than 20 years. However, while Health Authorities often generally supported 3Rs-aligned nonclinical programs, laws like the US’s Federal Food, Drug, and Cosmetic Act (FD&C Act) of 1938 still required that drugs be tested on animals before human trials. The FDA 2.0 Act of Dec 2022 removed the mandate for animal testing encouraging innovation in animal-free drug development. Furthermore, in April 2025, the FDA released a roadmap for new approach methodologies (NAMs), outlining the gradual phasing out of NHPs in preclinical safety studies beginning within the context of biologics’ safety assessment (FDA 2025). Similarly, the European Federation of Pharmaceutical Industries and Associations (EFPIA) has issued recommendations for phasing out animal testing. Their framework describes a continuum ranging from tests that can be discontinued immediately (where validated non-animal methods already exist) to areas where no alternatives are currently foreseeable, such as highly complex multi-generation studies, which lack viable *in vitro* or *in silico* models (EFPIA 2025).

In this evolving regulatory landscape, there is a growing need for human-relevant, mechanistically informative data sources that can support target qualification and safety assessment while reducing reliance on animal studies. In this context, ‘omics technologies are recognized as powerful tools to assess target expression across humans and preclinical species, and represent a critical step in drug R&D. It has been shown that expression levels of the targets are frequently considered when developing new chemical entities (69%) and always taken into account in the development of biologics (Namdari et al. 2021). This approach not only prioritizes promising targets but also flags potential adverse events, such as off-tumor activity, as seen in cardiopulmonary and skin toxicities with therapies targeting HER2 and EGFR, respectively (Tang et al. 2023). More recently, single-cell expression data have proven especially valuable for identifying cell type-specific markers, detecting low-frequency cell populations, cellular heterogeneity and cell-to-cell crosstalk (Chen et al. 2024).

Although numerous single-cell resources exist for other preclinical species (such as Tabula muris, which hosts multi-tissue single-cell datasets from mice), evaluating the expression of specific targets in the cynomolgus monkey model across multiple tissues remains challenging. Over the years, several organ-specific single-cell studies have been generated, but they differ considerably in methodologies, number of animals analyzed, tissue or anatomical coverage, granularity of cell type annotations, and even in the accessibility of their data (Peng et al. 2019; Wang et al. 2020, 2022; Zhou et al. 2024). Notably, two studies have reported multi-organ single-cell landscapes in cynomolgus monkeys (Han et al. 2022; Qu et al. 2022). Among these, Han et al. additionally provides harmonized cell labels across human and mouse datasets to enhance cross-species translatability. However, both studies are constrained by small number of individuals, and in the case of Han et al., the cross-species labeling does not encompass a broad range of organs.

This contrasts with the human context, wherein many single-cell landscapes have been released over the years. Notably, Tabula Sapiens provides annotations of millions of cells across multiple organs from healthy donors (The Tabula Sapiens Consortium* et al. 2022). Version 1 established a framework for addressing both technical and biological challenges, while version 2 expanded the resource to include 28 organs (up from 24) and incorporated data from multiple donors.

In recent years, cell type annotation has become increasingly accessible with the advent of new methodologies (Zeng et al. 2024; Cui et al. 2024). More recently, Rosen and collaborators (Rosen et al. 2023) have released the Universal Cell Embeddings (UCE) model, a foundation model trained on multi-species single-cell atlas data in a self-supervised manner without using cell type labels. UCE retains biological variation across tissues and species, is robust to technical noise and new cells from any species can be embedded into this space without additional model training, fine-tuning or labeling. This, in turn, enables an annotation by reference, *i.e.*, transferring labels from a well-annotated reference dataset to new datasets.

In this study, we aimed to expand the current single-cell transcriptomic landscape of *Macaca fascicularis* by integrating data from multiple studies at the organ level. To achieve consistent and cross-species annotation, we leveraged Tabula Sapiens V2 (comprising 28 organs as of December 2025) as the reference and applied UCE to harmonize cell type labels. Furthermore, in cases where no human counterpart was available, we kept original cell annotations or transferred labels across NHP studies, thereby increasing the number of animals represented while harmonizing annotations.. Overall, our results indicate robust cell annotation and a good agreement in the expression of various targets in human and cynomolgus monkeys but also highlight cases that may indicate species-specific responses. The framework provided here offers deeper insight into gene expression similarities and differences across organs, at cell type level and between species. By enabling early, cell type-resolved evaluation of cross-species target biology, this framework supports informed decisions on when cynomolgus monkey studies are likely to be predictive, when they may be refined, and when alternative approaches may be more appropriate – aligning with emerging 3Rs and NAMs guidance in biologics development.

## Material and methods

### 1. Datasets

#### 1.1 NHP cynomolgus (Macaca fascicularis) single-cell datasets

Single-cell RNA datasets were queried based on publications or curated databases such as GEO and ArrayExpress. Additional data (such as metadata and other annotations) were downloaed from Zenodo, such as in the case of Qu et al. Datasets were added based on queries up to January 2025. To be included in the study, the dataset was (i) explicitly indicated as sourced from the *M. fascicularis* species; (ii) a minimal set of files needed to be available for download (raw count matrices and feature or barcode cell annotation); (iii) if the study was tagged as case-control (*i.e.,* one group had undergone any sort of treatment or procedure), only control group individuals were used in downstream analyses. Count matrices and metadata from each study were then downloaded and converted into .h5ad objects as used by AnnData (v0.11.1) in Scanpy (Wolf et al. 2018). Each study was then individually assessed for quality metrics – since datasets were already processed and filtered by the authors, no additional filtering was applied except for the removal of cells with a mitochondrial gene expression percentage exceeding 35%.

For studies where only raw sequencing files were available, data were downloaded and processed using CellRanger (v8.0.0, 10x Genomics) and the genome annotated with the m.fascicularis v5 genome annotation. SoupX (Young and Behjati 2020) was applied to each experiment to correct for ambient RNA contamination. Corrected count matrices were then subjected to MADS (Median absolute deviations)-based filtering by removing cells with less than 200 detected genes and 500 total counts. Cells with mitochondrial gene expression above 5 MADs, or with ribosomal or hemoglobin expression above 5 MADs from the median were filtered out. Additionally, total counts and number of detected genes were required to fall between the 1st and 99.5th percentiles, removing extreme outliers in sequencing depth and complexity. Genes expressed in fewer than 3 cells were discarded. Scrublet (Wolock et al. 2019) was applied to identify potential doublets, which were removed based on a predicted doublet score using an expected doublet rate of 6%.

An overview of studies included (references and accession numbers) can be found in Supplementary Table S1.

#### 1.2. Human single-cell data

Human single-cell RNA data were retrieved from the Tabula Sapiens Consortium dataset (version 2) by downloading 28 single-cell datasets (per organ) from CELLxGENE (cellxgene.cziscience.com/collections/e5f58829-1a66-40b5-a624-9046778e74f5) (The Tabula Sapiens Consortium* et al. 2022).

### 2. Data analysis workflow

The analysis workflow was designed to enable systematic, scalable cross-species comparison of target expression at single-cell resolution, with the goal of informing translational safety assessment and reducing reliance on *in vivo* cynomolgus studies.

An overview of the analysis workflow is depicted in Suppl. Figure S1. Although this study primarily aimed to apply human-derived cell type annotations to the cynomolgus monkey datasets, some organs lacked a human counterpart set and therefore, in these specific cases, labels were transferred from any other cynomolgus monkey study that had cell annotations available. The study with the largest number of annotated cells across different tissues was used as the reference set for annotating other studies containing the same tissue or organ. An overview of label transfer direction, indicating which study served as the source for label transfer, is provided in Supplementary Table S1. In all cases, whenever original cell annotations were present, these were retained in the single-cell object.

#### 2.1. Universal cell embeddings (UCE)

Data integration using UCE was performed with the 4-layer model on the filtered, unnormalized count matrix for both NHP and human datasets. Uncorrected counts were always used as input, except for the CNP0001469 dataset, which only had SoupX (*32*) corrected counts for ambient RNA available. The “--species” flag was set to ‘macaca_fascicularis’ for NHP data and ‘human’ for human data. The resulting 1280-dimensional output matrix was stored as a new layer in the original AnnData object. No additional filtering or ortholog-based restrictions were applied.

#### 2.2 Cell label prediction

To predict and reannotate cell types in the NHP datasets, a k-nearest neighbors (KNN) model from the scikit-learn package (v1.3.0) (*33*) was trained, with *k* set to 20 neighbors and ‘distance’ as the weighting metric. Predictions were performed for each study and grouped at organ-level to avoid potential mislabeling caused by cells from unrelated organs exhibiting similar transcriptional profiles due to shared developmental lineage. The training dataset consisted of the UCE output from the human data or the cynomolgus monkey study used as reference for label transfer (Supplementary Table S1).

After training, the model was applied to predict cell types in the second, unlabeled dataset, using the respective UCE embeddings. Cell type predictions were retained only if the posterior probability of the top predicted cell type was 0.5 or higher; else cells were relabeled as “undefined”. Studies corresponding to the same organ were curated by evaluating organ-specific marker expression to ensure appropriate label transfer. During this process, studies in which marker genes showed widespread, non-specific expression across all cell types were excluded from the final dataset, as this pattern indicated insufficient data quality for inclusion in the atlas.

Across the integrated landscape, 27 organs were annotated using human reference labels. For the remaining organs, cell identities were either retained from the original publications or inferred through label transfer from a second cynomolgus study. Supplementary Table S1 provides an overview of all datasets and the corresponding source of cell labels.

#### 2.3 Study integration and ontology mapping

Following the analyses, all studies were consolidated into a single object, and the metadata were harmonized to ensure consistent column names and annotations across datasets. Raw gene expression was normalized for sequencing depth by scaling the total counts in each cell to 10,000, thereby enabling comparison of relative gene expression levels across cells. The normalized values were subsequently log-transformed. Missing information, such as sample format (cells or nuclei), sex, and age of the animals was retrieved by reviewing the original publications. Where available, animals were also assigned to age groups based on predefined intervals: infant (<=18 months), juvenile (>18 months – 4 years), young adult (> 4 –9.5 years), adult (>9.5 – 20 years) and old (>20 years) (Lau et al. 2020; Amato et al. 2022).

Following the integration, each study was annotated using an ontology-based approach, classifying samples according to anatomical location (e.g., duodenum, colon), organ (e.g., intestine), and biological system (e.g., gastrointestinal). When a study did not specify the exact anatomical region, the entire organ was used as the primary ontology level and maintained as a separate entity.

#### 2.4 Expression of organ-specific markers and use cases

A matrix containing tissue-enriched gene IDs across human organs was downloaded from the Human Protein Atlas (HPA v.25.0, updated November 11, 2025) (Thul and Lindskog 2018) and used to investigate organ-specific gene expression.

For each use case described below, expression data were retrieved from the full object to get organ/system-specific cells; percentage expression and average expression were calculated within each species and set using the list of genes retrieved from HPA and system-level ontologies (Supplementary Table S2). Genes were selected based on ortholog mapping between species. Plots were generated using Seurat (Stuart et al. 2019) and ggplot2 (Wickham 2009).

Subtypes and states of T cells were annotated using the STCAT tool (Shen et al. 2025). The PBMC set from NHPs (CNP0001469, 18,353 cells) and the Blood set from human (Tabula Sapiens V2, 85,233 cells) were used as inputs after rounding corrected counts to the nearest integer.

Drug-level information related to reported AEs were obtained from current FDA labels and links to documents mentioned in this study are detailed in Supplementary Table S3.

## Results

### 1. Landscape composition, cell labeling and expression of tissue-specific markers

After inclusion and analysis of all eligible datasets, the final merged dataset comprised 2,597,092 cells across 30 unique studies and 133 samples representing 57 anatomical locations, 43 organs, and 14 systems. An UMAP of the UCE embeddings across all studies is depicted in Figure 1A and the number of cells per anatomical site is depicted in Figure 1B.

**Figure 1.**
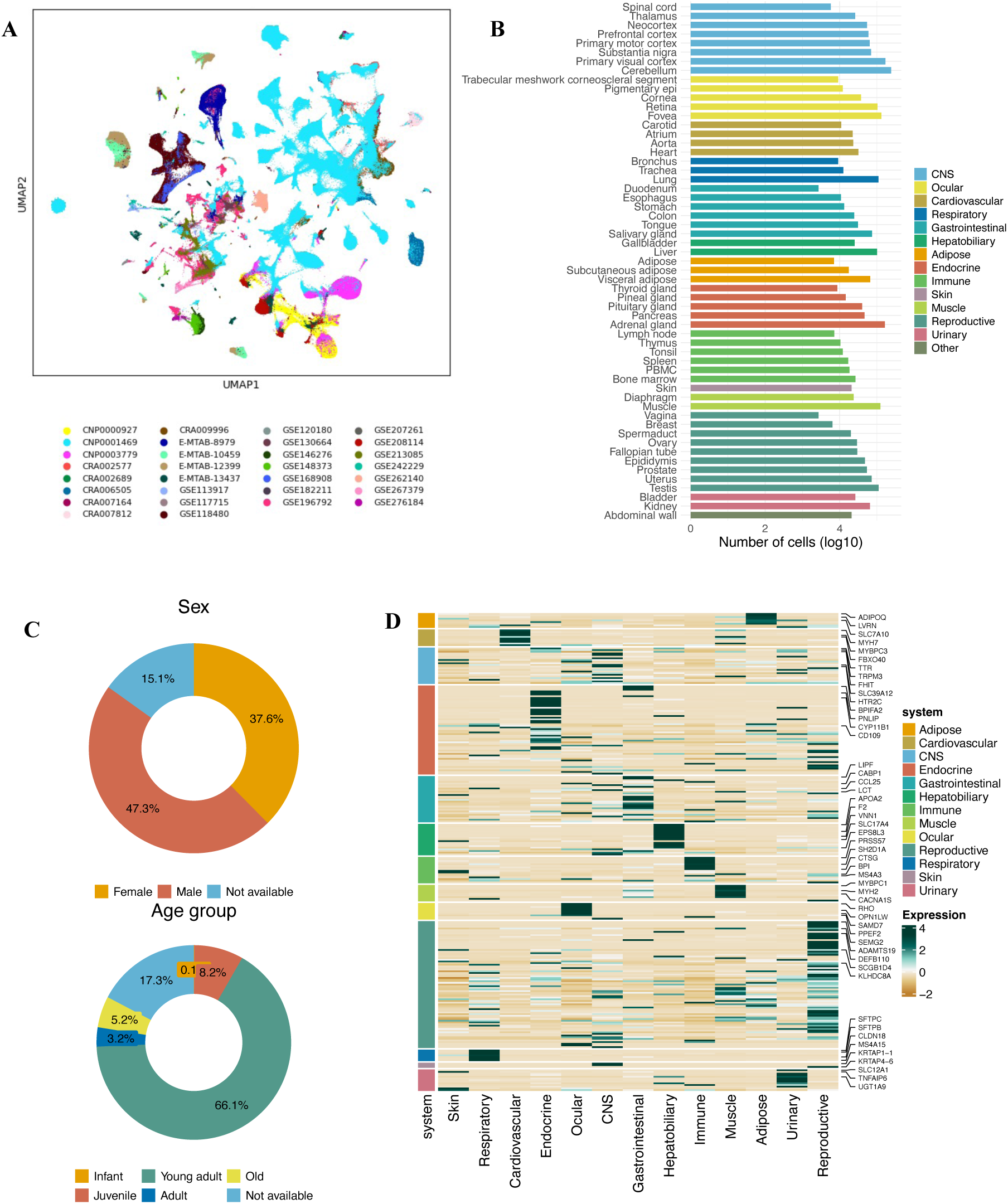
Non-human primate (NHP) *M. fascicularis* single-cell atlas composition and tissue-specific expression. (A) UMAP visualization of all cells in the atlas, colored by study of origin. The embedding was generated using the top UCE embeddings, with 30 nearest neighbors and cosine distance as the similarity metric. (B) Number of cells (log10 scale) across different anatomical parts, categorized by major organ systems. (C) Demographics of specimens included in this study, including sex (upper panel) and age group (lower panel). (D) Heatmap depicting the expression of human tissue-specific markers in cynomolgus organs spread across 13 systems.

Most cells originated from male subjects (47.3%), while 37.6% were from female subjects, and approximately 15% had unspecified sex. In terms of age classification, approximately 8% of cells were derived from juveniles, 66% from young adults, 3% from adults and 5% from aged animals. Less than 1% of all cells in the set originated from infants, while age group was not available for approximately 17% of the cells (see Figure 1C).

When evaluating cell annotation transfer, some studies exhibited greater variability across organs than others, with median KNN posterior probabilities ranging from 0.6 to 1 (Supplementary Figure S2). Despite this variability, most studies demonstrated high prediction confidence, and no major differences were observed between human-to-NHP (median = 0.95) and NHP-to-NHP (median = 1) posterior probability medians. A summary of the median posterior probabilities for labels transferred from Tabula Sapiens V2 is provided in Supplementary Figure S3. To further confirm that transferred labels reflected the biological identities of the cells, we performed comparisons between studies whose original cell annotations were available across multiple tissues.

For human-to-NHP evaluation, we compared original labels provided by Han et al. (2022) and Tabula Sapiens V2 final predicted cell types. Sankey plots for each organ indicated a good agreement between major cell type annotations, with some minor differences (Supplementary Figure S4). While large cell populations generally retained consistent identities across datasets, some organs reveal differences in annotation granularity. For example, in the bladder, the original dataset (Han et al., 2022) includes finer distinctions such as “cycling intermediate,” “intermediate,” “umbrella,” and “urothelial” cells, whereas the transferred annotations are consolidated under “bladder urothelial cell,” the annotation in the reference set available for specialized bladder cells. Concerning label transfers between cynomolgus monkey studies, a higher variability in the quality of labels transfers were observed, in which original labels from Qu et al. (2022), were compared to labels transferred from the Han et al. (2022) as an example (Supplementary Figure S5). Organs with lower median prediction probabilities (e.g., trachea and uterus) showed a higher proportion of undefined cells. In other cases, median probabilities were higher overall, yet label discordance was observed. For example, in aorta, cells originally annotated as ‘smooth muscle cells’ were reassigned as ‘endothelial cells’ following label transfer. These discrepancies may reflect technical factors such as differences in sample preparation and preprocessing, which can increase doublet rates or ambient RNA contamination, as well as QC filtering criteria that may preferentially retain certain cell types. They may also arise from domain shift between query and reference datasets, from inconsistencies or potential mislabeling in either the reference or query labels, or from incomplete representation of organ-specific or germline cell states in the training data. These factors may lead to reduced confidence or biologically implausible nearest-neighbor assignments; inspection of cluster structure, application of more stringent KNN probability cutoffs, and review of cell-specific marker expression by domain experts can help mitigate such implausible annotations.

Across all systems, expression of human-based, tissue-specific targets showed consistent clustering in the cynomolgus monkey landscape, indicating overall similar expression of the main markers (Figure 1D). Together, these results establish a robust foundation for downstream translational analyses by ensuring that observed expression differences reflect biological rather than technical variation.

### 2. Use cases

To illustrate how this atlas can inform translational risk assessment and support refinement or reduction of NHP studies, we present a series of use cases relevant to biologics development.

#### 2.1. Expression of targets with translationally concordant adverse events and matched tissue-level expression

Publicly available databases were systematically interrogated to identify marketed antibodies and antibody-drug conjugates that showed concordant, target-attributed toxicity in humans and cynomolgus monkeys affecting the same tissue or organ. *CD44, CD47, EGFR, F3, NECTIN4* and *VEGFR2* were selected as candidates. Current knowledge regarding their expression in both species and adverse events described in preclinical studies and clinical findings are described in Table 1.

**Table 1.**
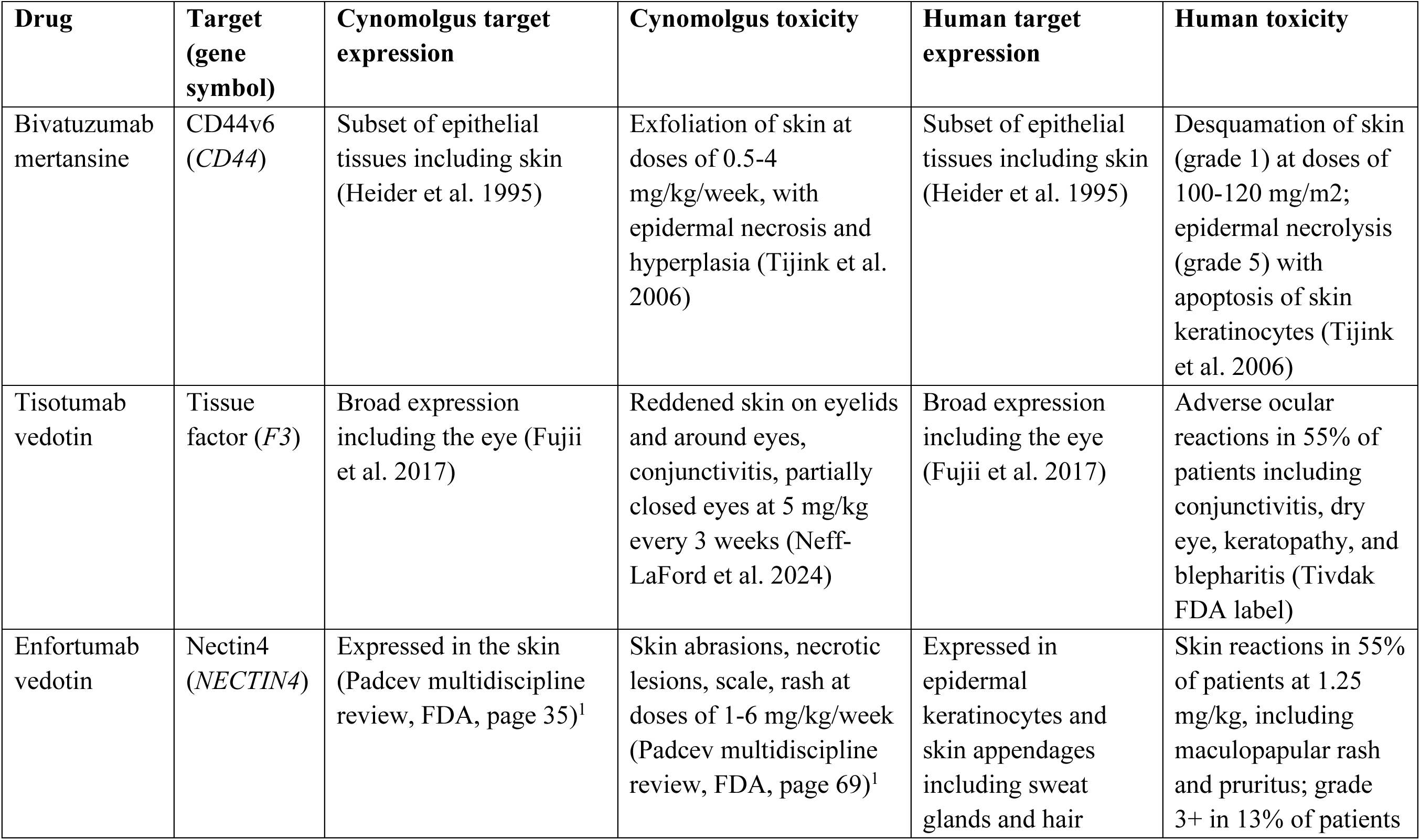
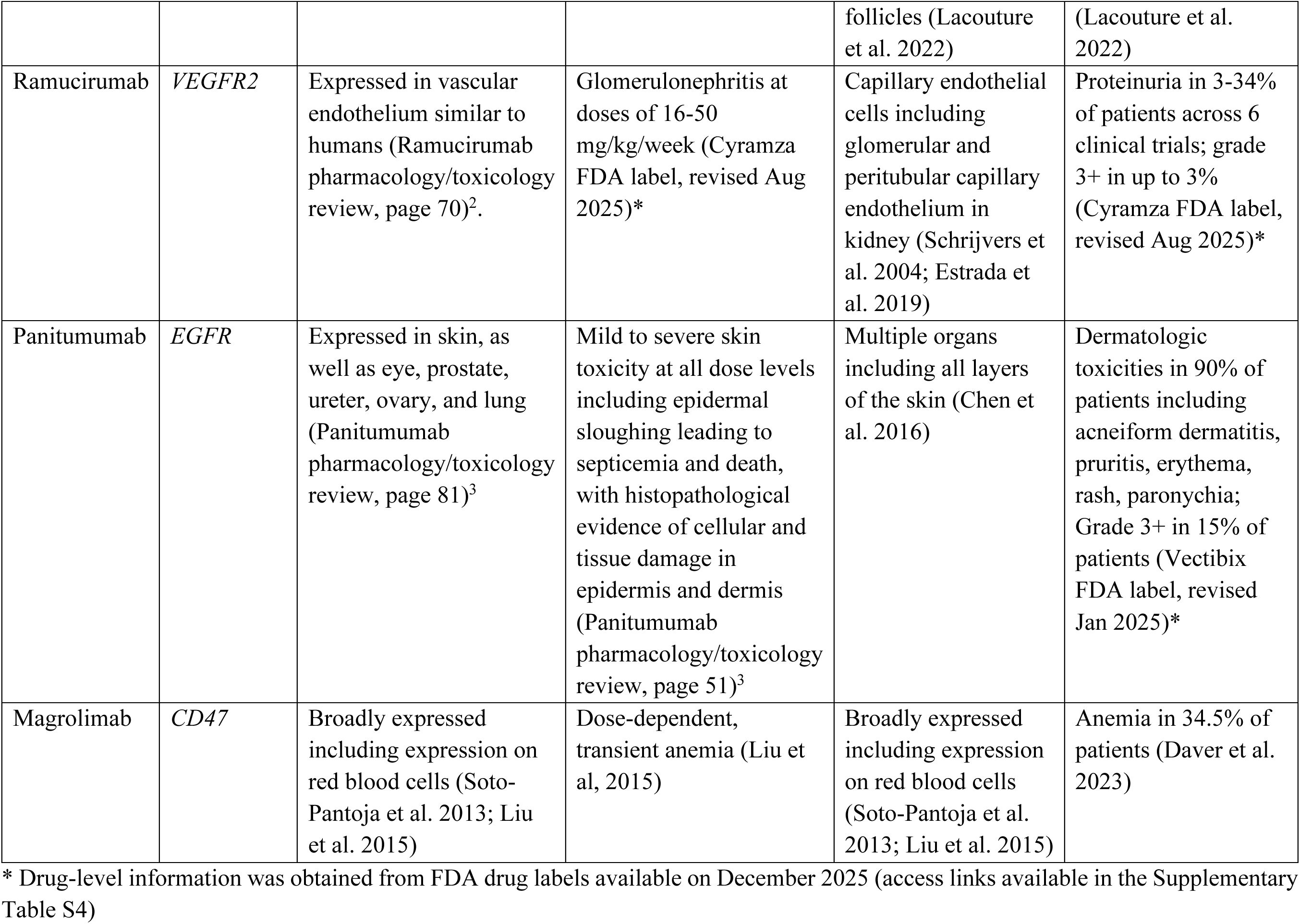
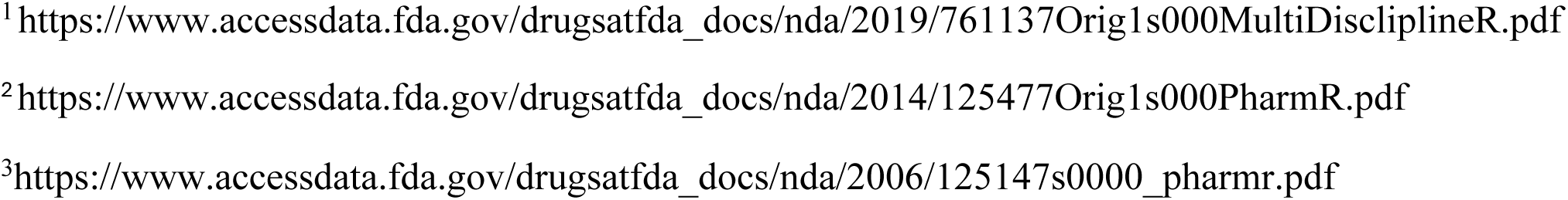
Antibodies and ADCs with concordant target expression and toxicity in humans and cynomolgus monkeys.

| Drug | Target<br>(gene<br>symbol) | Cynomolgus target<br>expression | Cynomolgus toxicity | Human target<br>expression | Human toxicity |
| --- | --- | --- | --- | --- | --- |
| Bivatuzumab<br>mertansine | CD44v6<br>(*CD44*) | Subset of epithelial<br>tissues including skin<br>(Heider et al. 1995) | Exfoliation of skin at<br>doses of 0.5-4<br>mg/kg/week, with<br>epidermal necrosis and<br>hyperplasia (Tijink et al.<br>2006) | Subset of epithelial<br>tissues including skin<br>(Heider et al. 1995) | Desquamation of skin<br>(grade 1) at doses of<br>100-120 mg/m<sup>2</sup>;<br>epidermal necrolysis<br>(grade 5) with<br>apoptosis of skin<br>keratinocytes (Tijink<br>et al. 2006) |
| Tisotumab<br>vedotin | Tissue<br>factor (*F3*) | Broad expression<br>including the eye (Fujii<br>et al. 2017) | Reddened skin on eyelids<br>and around eyes,<br>conjunctivitis, partially<br>closed eyes at 5 mg/kg<br>every 3 weeks (Neff-LaFord et al. 2024) | Broad expression<br>including the eye<br>(Fujii et al. 2017) | Adverse ocular<br>reactions in 55% of<br>patients including<br>conjunctivitis, dry<br>eye, keratopathy, and<br>blepharitis (Tivdak<br>FDA label) |
| Enfortumab<br>vedotin | Nectin4<br>(*NECTIN4*) | Expressed in the skin<br>(Padcev multidiscipline<br>review, FDA, page 35)<sup>1</sup> | Skin abrasions, necrotic<br>lesions, scale, rash at<br>doses of 1-6 mg/kg/week<br>(Padcev multidiscipline<br>review, FDA, page 69)<sup>1</sup> | Expressed in<br>epidermal<br>keratinocytes and<br>skin appendages<br>including sweat<br>glands and hair | Skin reactions in 55%<br>of patients at 1.25<br>mg/kg, including<br>maculopapular rash<br>and pruritus; grade<br>3+ in 13% of patients |
|  |  |  |  | follicles (Lacouture et al. 2022) | (Lacouture et al. 2022) |
| Ramucirumab | *VEGFR2* | Expressed in vascular endothelium similar to humans (Ramucirumab pharmacology/toxicology review, page 70)<sup>2</sup>. | Glomerulonephritis at doses of 16-50 mg/kg/week (Cyramza FDA label, revised Aug 2025)* | Capillary endothelial cells including glomerular and peritubular capillary endothelium in kidney (Schrijvers et al. 2004; Estrada et al. 2019) | Proteinuria in 3-34% of patients across 6 clinical trials; grade 3+ in up to 3% (Cyramza FDA label, revised Aug 2025)* |
| Panitumumab | *EGFR* | Expressed in skin, as well as eye, prostate, ureter, ovary, and lung (Panitumumab pharmacology/toxicology review, page 81)<sup>3</sup> | Mild to severe skin toxicity at all dose levels including epidermal sloughing leading to septicemia and death, with histopathological evidence of cellular and tissue damage in epidermis and dermis (Panitumumab pharmacology/toxicology review, page 51)<sup>3</sup> | Multiple organs including all layers of the skin (Chen et al. 2016) | Dermatologic toxicities in 90% of patients including acneiform dermatitis, pruritis, erythema, rash, paronychia; Grade 3+ in 15% of patients (Vectibix FDA label, revised Jan 2025)* |
| Magrolimab | *CD47* | Broadly expressed including expression on red blood cells (Soto-Pantoja et al. 2013; Liu et al. 2015) | Dose-dependent, transient anemia (Liu et al, 2015) | Broadly expressed including expression on red blood cells (Soto-Pantoja et al. 2013; Liu et al. 2015) | Anemia in 34.5% of patients (Daver et al. 2023) |

\* Drug-level information was obtained from FDA drug labels available on December 2025 (access links available in the Supplementary Table S4)
<sup>1</sup>https://www.accessdata.fda.gov/drugsatfda_docs/nda/2019/761137Orig1s000MultiDiscliplineR.pdf
<sup>2</sup>https://www.accessdata.fda.gov/drugsatfda_docs/nda/2014/125477Orig1s000PharmR.pdf
<sup>3</sup>https://www.accessdata.fda.gov/drugsatfda_docs/nda/2006/125147s0000_pharmr.pdf

Figure 2A shows a comparison of the ranked, pseudobulk expression of these targets in human and cynomolgus monkey datasets across each target, indicating that, aside from vascular endothelial growth factor-2 (*VEGFR2*, also known as kinase insert domain receptor), most targets show good correlation when comparing average expression across all organs (Spearman’s rho range 0.32-0.59).

**Figure 2.**
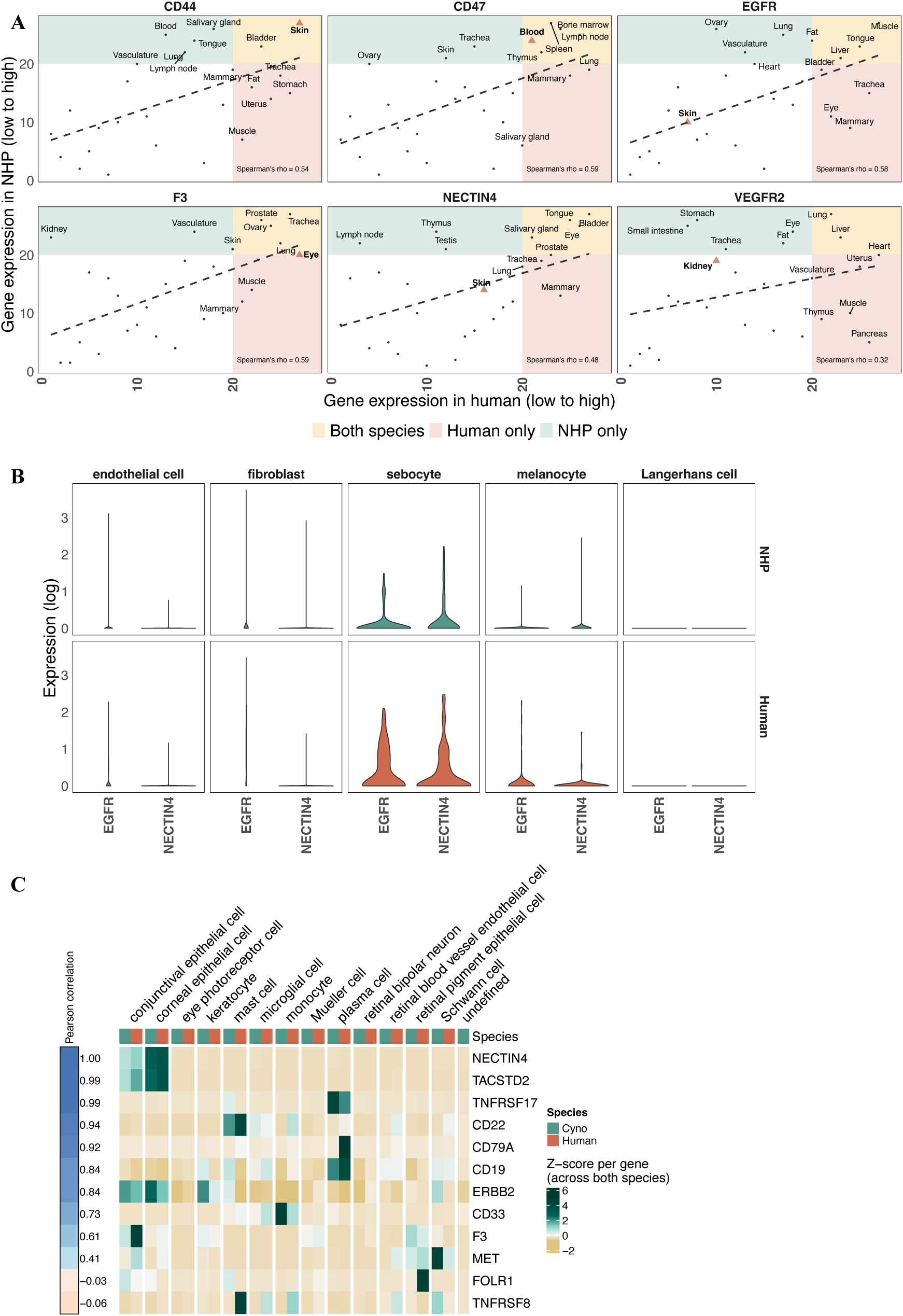
Cross-species comparison of biologics’ targets across multiple organs and specific cell types in skin and eye. **(A)** Rank-rank plots for each target, indicating organ-level expression ranks in human versus the non-human primate (NHP) cynomolgus monkey. Twenty-seven tissues common to Tabula Sapiens V2 and the cynomolgus single-cell atlas generated in this study are shown in each plot. Expression ranks were calculated independently within each species based on organ-level expression values, with higher ranks indicating higher expression. Triangles indicate organs corresponding to reported adverse event sites. Shaded regions denote concordant or discordant expression ranking between species: high expression in both species (yellow), high in human and low in cynomolgus monkey (red), or high in cynomolgus monkey and low in human (green). Dashed lines represent the Spearman rank correlation across organs, with the correlation coefficient shown in each panel. **(B)** *EGFR* and *NECTIN4* expression in cynomolgus and human sets across cell types in skin. Labels are harmonized between species following UCE and cell type prediction with KNN. Expression values are log-transformed. **(C)** Expression of ADC targets across different cell types in the eye. Labels are harmonized between species following UCE and cell type prediction with KNN. Pearson correlation coefficients were calculated on log-transformed expression values, while scaling (z-score transformation) across species and genes was performed for visualization.

Across all targets, a median of approximately 53% of top-ranked organs (rank ≥20) were shared between human and cynomolgus monkey, where overlap was calculated as the percentage of organs ranked ≥20 in both species relative to the total number of high-ranked organs per target (range 25%-75%). *CD44* showed high expression in skin in both human and cynomolgus monkey, while *F3* (also known as tissue factor) demonstrated expression in ocular tissues in both species. A similar pattern was observed for *CD47*, with high-ranking expression in blood in both human and cynomolgus monkey. For *VEGFR2*, expression was detectable in endothelial-rich tissues across species, although kidney ranks differed between them.

*EGFR* exhibited strong expression across multiple tissues while *NECTIN4* was moderate and restricted to a few sites; in both cases, however, the tissues associated with their reported adverse events (i.e., skin) did not appear among the top-ranked sites. Examination of single-cell-level expression of *EGFR* and *NECTIN4* showed they are prominently expressed in sebocytes, melanocytes and keratinocyte cell types (Figure 2B).

By identifying organs and cell types with conserved, target-level expression across species – and distinguishing these from species– or tissue-specific differences – this landscape enables prospective selection of relevant tissues for focused safety assessment in cynomolgus monkey. This analysis establishes a baseline expectation for translational concordance, supporting the use of targeted, hypothesis-driven NHP studies rather than broad exploratory programs.

#### 2.2. Ocular toxicity and antibody-drug conjugates (ADC)

Within the realm of biologics, antibody-drug conjugates (ADCs) represent a class of immunoconjugates in which a monoclonal antibody (mAb) is covalently linked to a cytotoxic payload. As of December 2025, 15 ADCs have been approved for diverse oncological indications by the FDA. A description of these drugs, their targets and payload (classes) are described in Table 2.

**Table 2.**
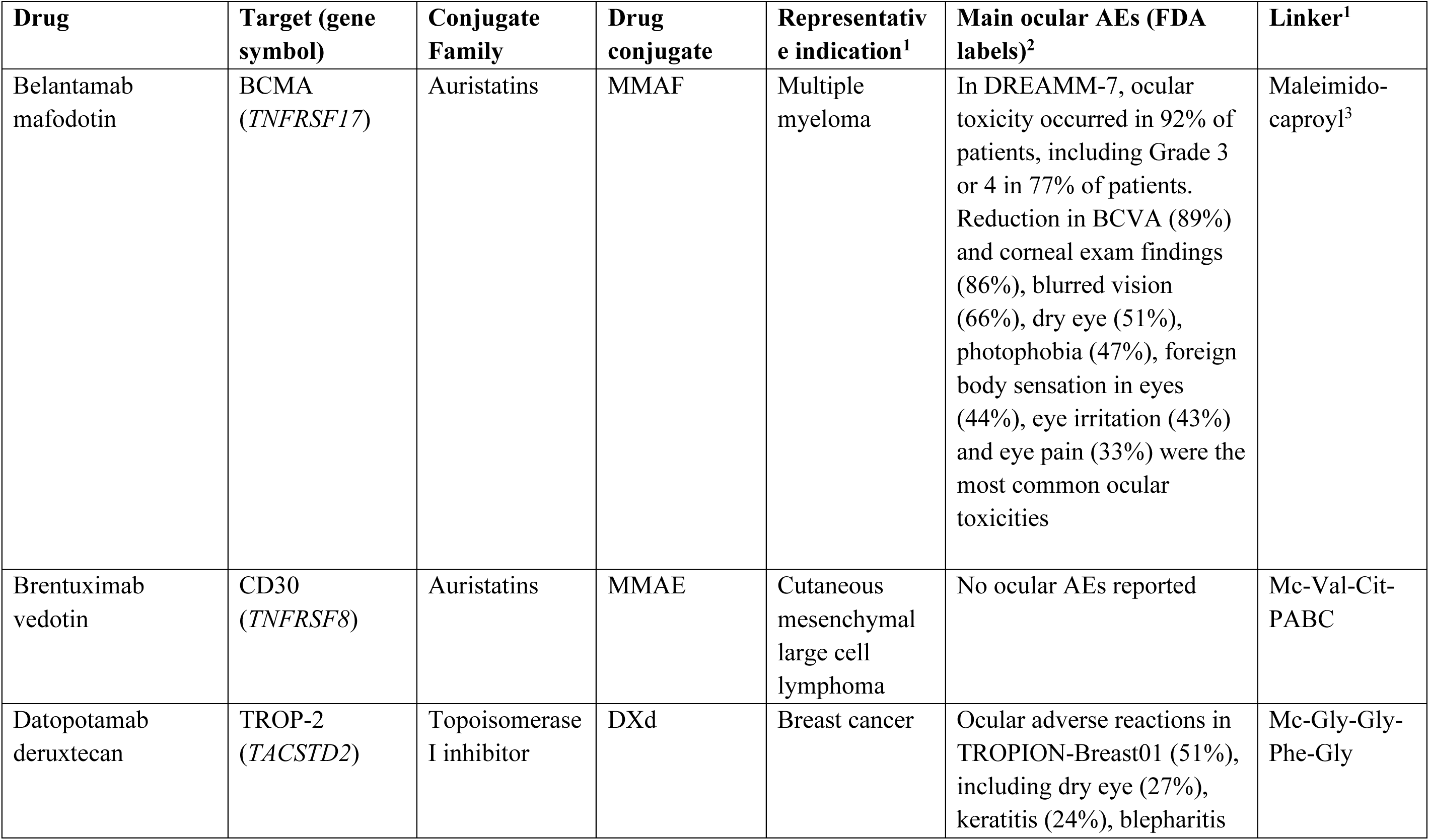
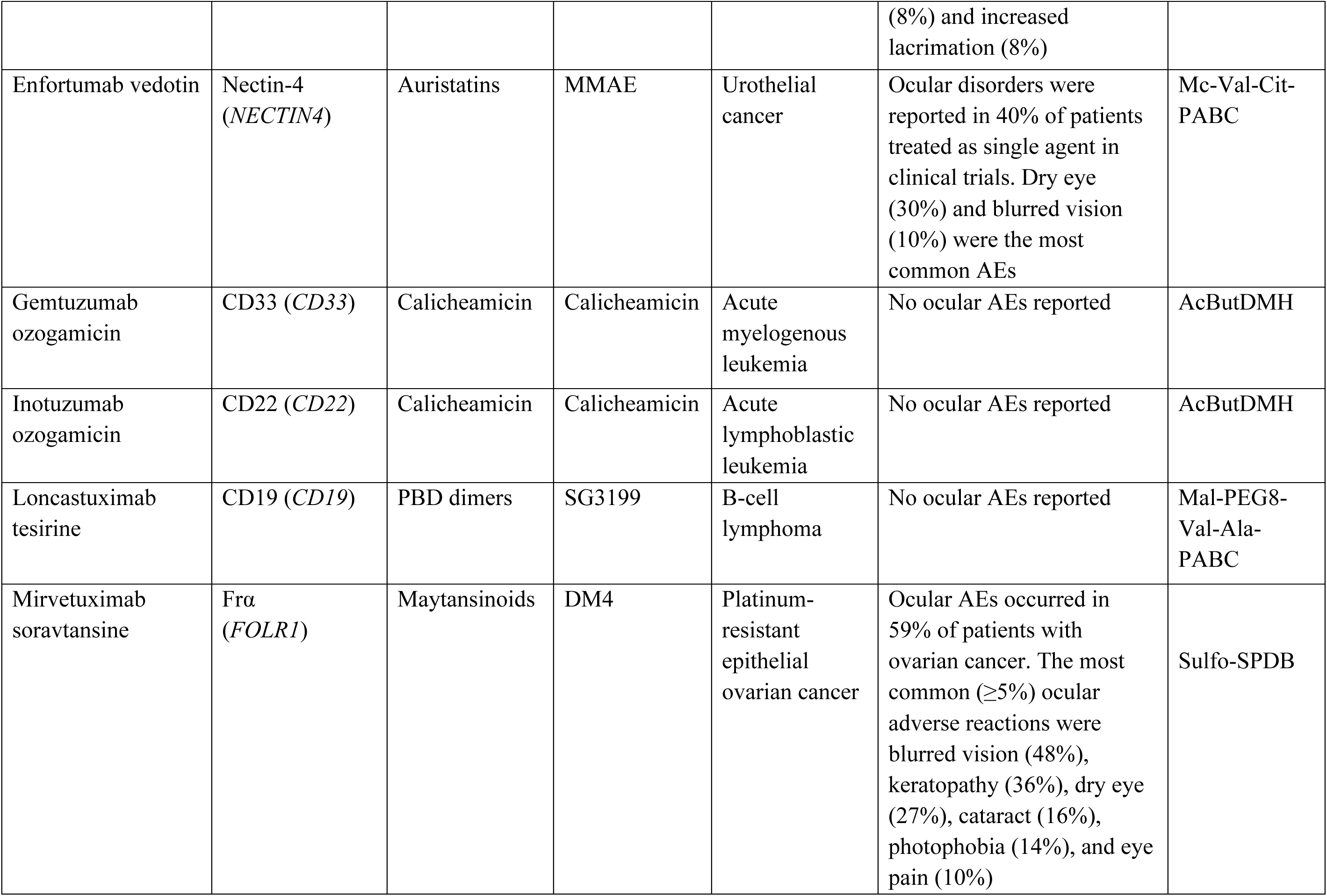
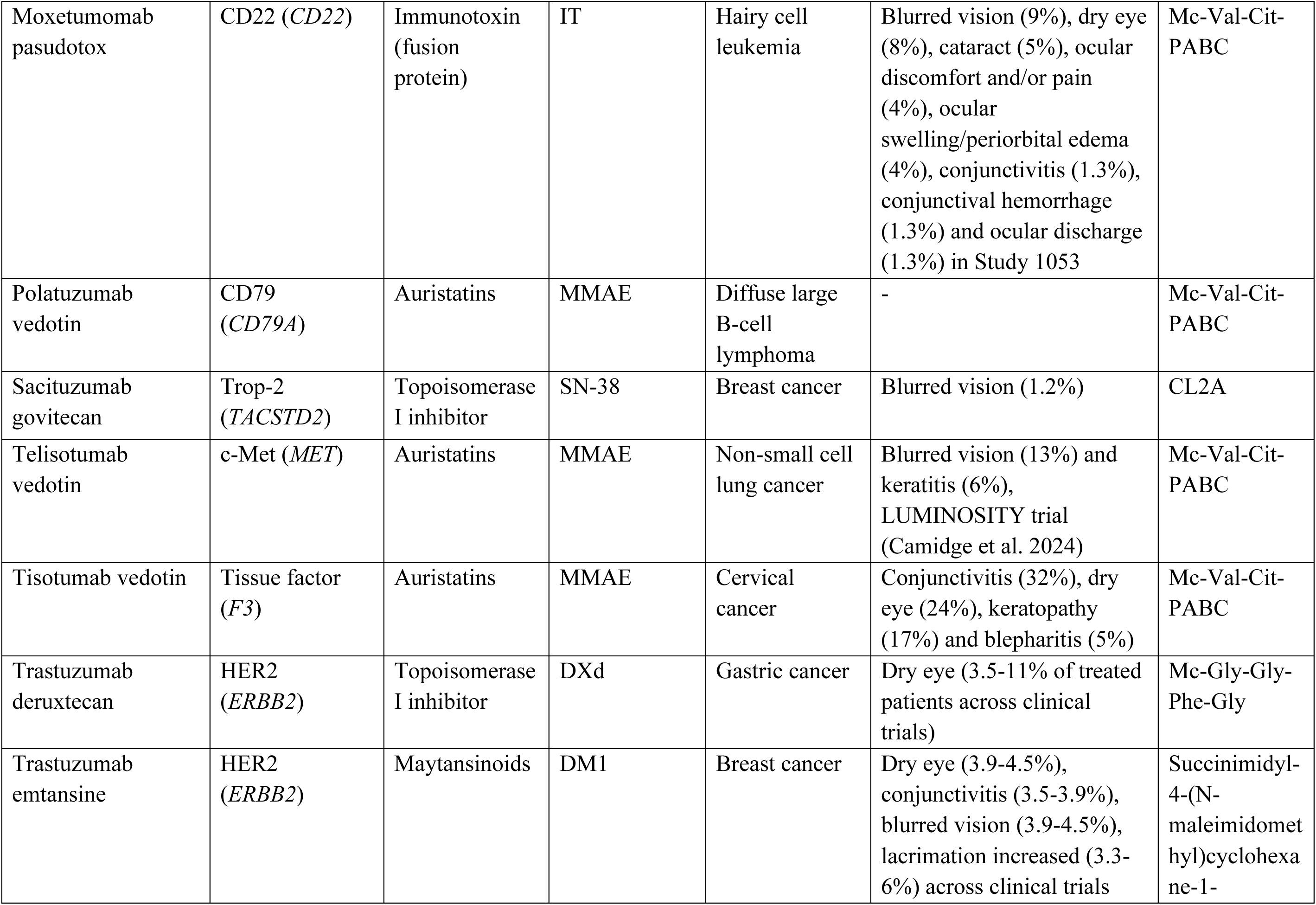
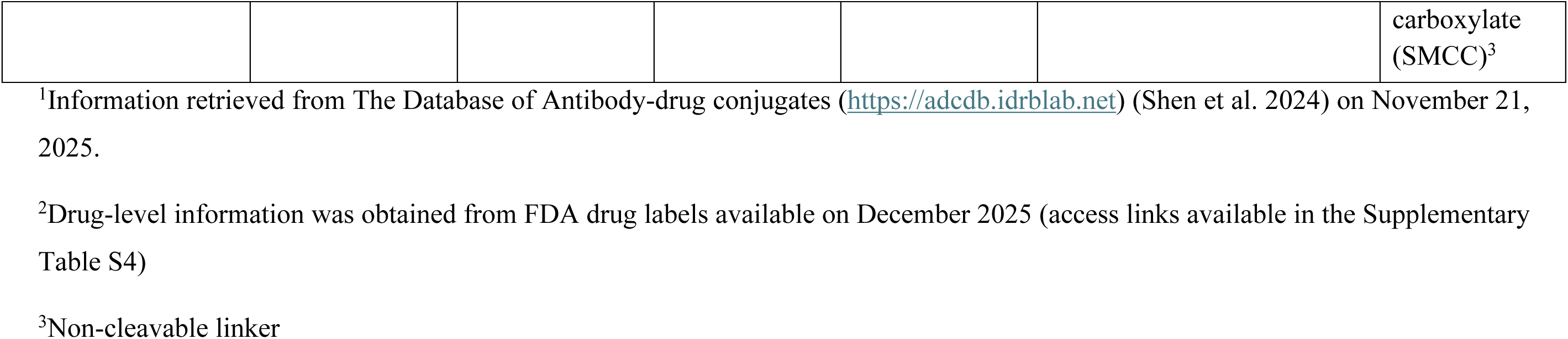
List of FDA-approved ADCs, their composition, oncological indications and ocular AEs associated with treatment.

Among others, ocular toxicities are one of the main AEs associated with ADC treatment, and the exact underlying mechanisms remain poorly understood. Investigating target expression in specific compartments of the eye may be helpful to estimate whether the observed AEs can be (partially) explained by on-target, off-site binding.

Expression of ADC targets was observed across cells in distinct parts of the eye and in multiple cell types (Figure 2C). For each gene, log-normalized average expression values were calculated per matched cell type in each species. Pearson correlation coefficients were then computed between human and NHP across matched cell types to quantify conservation of cell-type-specific expression patterns. No scaling was applied prior to correlation analysis. For visualization purposes only, gene expression values were standardized to z-scores per gene across the combined dataset (both species and all cell types) to harmonize dynamic range across genes. Overall, the matrix indicates varying degrees of conservation across genes and cell types. *NECTIN4*, *TACSTD2*, *TNFRSF17*, *CD22*, *CD79A*, *CD19*, and *ERBB2* showed strong concordance between human and cynomolgus z-scores (Pearson correlation > 0.80), while *CD33* and *F3* displayed moderate correlation (Pearson correlation 0.73 and 0.61, respectively).

*NECTIN4*, *TACSTD2*, and, to a lesser extent, *ERBB2* demonstrated concordant expression in conjunctival and epithelial cells in both human and cynomolgus. In addition, *TNFRSF17*, *CD79A*, and *CD19* – targets of ADCs used in hematological indications – showed comparable expression rankings between human and NHP, particularly in plasma cells. Notably, *FOLR1* and *TNFRSF8* exhibited little to no concordance in expression rank across eye cell populations between human and NHP (Pearson correlation –0.03 and –0.06, respectively). Both genes showed marked species-specific patterns, with high relative expression in certain cell types in one species but not the other. For example, *FOLR1* rank was high in conjunctival epithelial cells in cynomolgus (albeit with low overall expression) but not in human, whereas the opposite pattern was observed in retinal pigment epithelial cells, where expression rank was high in human but low in cynomolgus.

By resolving target expression at the level of specific ocular cell populations, this analysis achieves anatomical and cellular resolution not attainable with bulk tissue profiling and further uncovers interspecies differences. These results illustrate how cell-resolved, *in silico* analyses can serve as an upstream decision-support step, providing anatomical context that guides and complements subsequent *in vivo* experimentation.

#### 2.3 CD28 expression across T-cell subpopulation and states

Given the central role of CD28 in T-cell stimulation and its implication in one of the most severe immune-related clinical trial failures, we specifically examined CD28 expression patterns across species to assess whether our approach could recapitulate known cell-specific expression differences (Eastwood et al. 2010; Hünig 2016). Blood-derived samples from human and non-human primates originated from whole blood and PBMC fractions, respectively. STCAT was applied to both sets: 64% of cells (11,844) in the PBMC set in cynomolgus monkey were identified as T-cells, in contrast to 11% of cells derived from human blood (9,787) (Figure 3A). Despite this compositional difference, the comparison remains appropriate for exploratory purposes, as both datasets capture overlapping immune cell compartments and allow assessment of cross-species label transfer performance within shared cell types. Labels transferred from human to cynomolgus monkey indicated a high degree of concordance between predicted cell types (Tabula Sapiens V2 following UCE) and original cell annotations as defined by Han et al., 2022 (Figure 3B).

**Figure 3.**
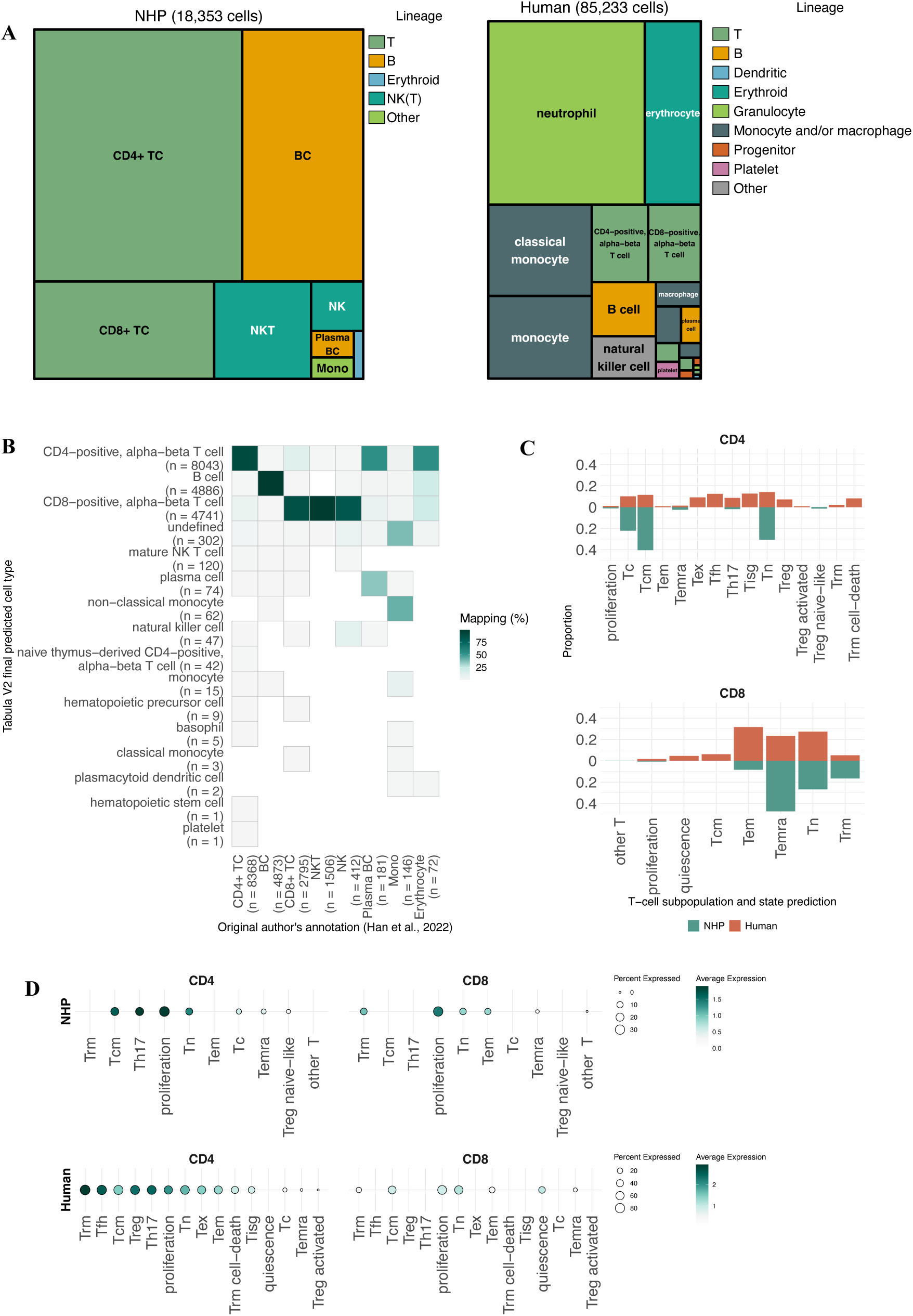
T-cell states and *CD28* expression across cell populations in human and cynomolgus monkey. **(A)** Cell type composition in single-cell sets obtained from the non-human primate (NHP) cynomolgus monkey (PBMC, left panel) and human (Blood, right panel). Cells were color-coded based on lineage classification. **(B)** Comparison between the original author’s annotation (x-axis, Han et al., 2020) and predicted Tabula Sapiens V2 cell labels (y-axis), represented as the percentage of cells predicted in each category. **(C)** Prediction proportion for CD4 and CD8 T-cell states in human and cynomolgus predicted by STCAT. **(D)** CD28 expression in CD4 and CD8 T-cell states in humans and in the NHP cynomolgus monkey. Expression (log-transformed) was calculated within each set (human or cynomolgus).

After assignment of T-cell subpopulation/states, results indicated that, despite the lower number of T-cells (both absolute and relative to the total number of cells), the human set showed a higher diversity of subtypes (Figure 3C). Similarly, expression of CD28 was observed to be moderate to high across subsets of cell populations in cynomolgus monkey, but overall, more pronounced in the human set, especially among CD4 subtypes (Figure 3D). This case illustrates how the atlas not only identifies conserved biology but also flags critical species-specific differences that may contraindicate reliance on cynomolgus models.

## Discussion

We present a harmonized single-cell atlas of *Macaca fascicularis* that integrates publicly available datasets spanning 43 organs and 14 physiological systems. Using foundation-based cross-species annotation, we applied UCE to align cellular identities between cynomolgus monkey and human at an unprecedented scale and resolution. The resulting atlas, comprising over 2.5 million cells from 30 studies, demonstrates strong prediction probabilities and high concordance of cell type annotations in datasets with known labels. These results hold both within and across species, underscoring the robustness of our approach in capturing biological signals intrinsic to cell identity. Given that our annotation strategy leverages Tabula Sapiens V2 as the human reference atlas, our use cases focus on organs shared between the sets to best illustrate the practical application of the cross-species mapping strategy.

To evaluate translational relevance, we first identified biologics with concordant toxicity profiles in both human and cynomolgus monkey, *i.e*., where treatment with the same molecule results in adverse events (AEs) affecting the same tissues in both species. We then assessed whether the expression of biologics’ targets is similarly enriched in the affected tissue(s) across species. This concordance of response (toxicity) followed by supporting expression is not intended to imply a mechanistic causal relationship, but rather to evaluate whether target expression may contribute to, or help explain, observed tissue-specific AEs. *CD44* (skin), *CD47* (blood) and *F3* (eye) showed cross-species concordance, all figuring among top-ranking gene-AE site matches in human and cynomolgus monkey. *CD47* was particularly consistent across organs enriched with blood and immune cells (*i.e.,* blood, spleen, bone marrow, lymph node and thymus).

This is noteworthy considering that the main adverse events in anti-CD47 therapy are hematologic issues such as anemia and thrombocytopenia (Table 1); single-cell analyses further localized *CD47* expression to platelets in human and hematopoietic stem cells in cynomolgus (Supplementary Figure S7). Together, these results demonstrate how integrating organ-level toxicity concordance with cross-species, cell type-resolved expression data can support biologically informed assessment of translational risk and help rationalize the continued (or reduced) use of the cynomolgus model in preclinical safety evaluation.

To illustrate how cell type-specific expression improves the translational interpretation of target-associated toxicity, we next examined skin-related targets. Despite their well-documented cutaneous adverse events, *EGFR* and *NECTIN4* did not rank among the top gene-organ expression matches in either human or cynomolgus monkey, indicating that tissue-level expression metrics alone are insufficient to predict organ-specific toxicity. In contrast, single-cell analyses revealed marked enrichment of *EGFR* and *NECTIN4* within discrete epithelial subpopulations, most notably keratinocytes and sebocytes in both species.

These findings provide a mechanistic basis for the translational relevance of cynomolgus monkey as a preclinical model for skin toxicity, despite limited signal at the organ level. EGFR signaling is essential for epidermal homeostasis, with inhibition driving keratinocyte differentiation and compromising skin barrier integrity, as well as impairing sebocyte lipogenesis, consistent with acneiform lesions observed clinically following EGFR blockade (Dahlhoff et al. 2015; Joly-Tonetti et al. 2021; Bierbrier et al. 2023). Decreased EGFR signaling has been shown to directly impact lipogenesis in sebocytes, a key feature of sebaceous gland-associated skin diseases, such as acne (Dahlhoff et al. 2015). This is in line with some varieties of anti-EGFR skin injuries, which may manifest as acneiform lesions (Bierbrier et al. 2023). Notably, this restricted cellular expression pattern differs from that of *CD44*, which is broadly expressed across skin cell types and shows strong concordance at the organ level (Supplementary Figure S6). Collectively, these results demonstrate that integrating single-cell expression profiles with organ-level analyses may enable more accurate prediction of target-mediated toxicities and supports rational species selection in preclinical safety assessment.

This is particularly critical when evaluating tissues and organs that are difficult to obtain and characterize, such as ocular tissues. With the expanding number of AEs for approved ADCs, safety concerns have intensified, underscoring the need for robust alternative models and data that are relevant for the human context. AEs can largely vary across ADCs, ranging from mild complaints (*e.g.,* dry eyes, increased blinking or lacrimation) to severe outcomes (e.g., cataract, keratitis), indicating the involvement of different anatomical compartments. In this study, we take the first step toward assessing whether these effects may be due to potential on-target, off-site effects in specific cell populations in the eye. We show that while targets from hematological malignancies show localized expression in blood cell populations (*e.g., TNFRSF17* in plasma cells, *CD33* in monocytes and *CD22* in mast cells), targets from solid indications are shown to be expressed in conjunctival (*MET, F3, ERBB2, NECTIN4, TACSTD2*), corneal epithelial cells (*ERBB2*, *NECTIN4, TACSTD2*) and retinal pigment epithelial cells *(MET, FOLR1*) at various degrees.

Taken together, these data offer a compelling mechanistic rationale for the observed incidence of ocular AEs in patients receiving ADCs against solid tumor targets. For instance, *NECTIN4* is expressed in both conjunctival and corneal epithelial cells, and AE reports have indicated the occurrence of dry eyes and blurred vision (Table 2). Notably, *ERBB2*, the target of both trastuzumab deruxtecan (TD) and trastuzumab emtansine (TE), also appears in conjunctival and corneal epithelial cells, and AEs linked to these drugs include, at low frequencies (11% or less across treated patients), dry eye, conjunctivitis and increased lacrimation (Table 2). Notably, the cross-species component of the atlas enables detection of potential translational gaps, as exemplified by the divergent *FOLR1* and *MET* expression profiles between human and cynomolgus monkey eye tissues. Although it should be noted that other factors may influence ADC-induced ocular toxicity (*e.g.* payload type, linker, drug-to-antibody ratio, bioavailability within the eye), these results highlight how the atlas allows organ– and cell type-level insight even for tissues difficult to sample *in vivo*.

Furthermore, certain safety liabilities may only become apparent when examining gene expression at even higher resolution, down to the level of cell subtypes or activation states, which can only be investigated through probing robust markers in single cells. Theralizumab (TGN1412) was a CD28-targeting T-cell agonist developed for the purpose of balancing the immune system and counteracting autoimmune diseases such as rheumatoid arthritis (Eastwood et al. 2010; Hünig 2016). Despite showing no toxicity in rodent and NHP studies, the drug led to catastrophic outcomes in first-in-human trials, where all participants experienced severe systemic inflammatory responses. This case represents one of the relatively rare, but highly consequential, instances in which preclinical species fail to anticipate human-specific toxicities. Subsequent studies revealed that *CD28* is expressed in CD4+ effector memory T-cells of humans but not other species commonly used in preclinical assessments (Eastwood et al. 2010). By computationally predicting T-cell subpopulations across species, we demonstrate that humans exhibit greater T-cell heterogeneity based on canonical human markers, and crucially, confirm the expression of *CD28* in human CD4⁺ effector memory cells – which is not mirrored in cynomolgus monkey. These findings highlight how integrated cross-species immune profiling can uncover subtle yet critical differences that are often overlooked by traditional R&D pipelines.

Together, these findings demonstrate how a unified cynomolgus single-cell atlas can reveal conserved biology and translational potential, pinpoint species-specific differences, and improve interpretation of tissue– and cell type-specific toxicities. The use cases presented (spanning toxicity concordance, ocular liabilities, and immune profiling) collectively demonstrate that this atlas provides actionable translational insight not possible with existing fragmented NHP datasets. By enabling high-resolution, high-throughput cross-species comparisons, this resource strengthens target qualification, enhances mechanistic understanding, and supports a more predictive and ethically aligned path for biologics development. Rather than advocating wholesale replacement of NHP models, this framework enables evidence-based reduction and refinement by identifying both conserved and species-specific aspects of target biology, directly supporting the intent of recent FDA and EFPIA guidance on new approach methodologies.

## Conclusions

In this study, we demonstrate the feasibility and utility of integrating heterogeneous single-cell datasets from *Macaca fascicularis* by harmonizing cell annotations across studies, thereby significantly expanding the number of cells and subjects available for gene expression analysis. Our approach addresses a longstanding challenge in preclinical research: the fragmentation and variability of non-human primate (NHP) transcriptomic data. While human and murine single-cell atlases have matured into comprehensive resources, NHP datasets remain sparse, organ-specific, and in some cases inconsistently (and even sparsely) annotated. By implementing a scalable, model-based framework for cell label harmonization, we broaden the utility of existing NHP data and establish a methodology that is readily adaptable to other species and experimental contexts. We demonstrate, through different use cases in the field of biologics, how this single-cell atlas can be used to identify target expression at a single-cell resolution, as well as investigating cross-species differences.

Although this framework presents challenges related to the consolidation of various experimental designs (*i.e.,* tissue sample collection, biological fractions used, data preprocessing, metadata availability and annotation) as well as the substantial computational requirements, these can be manageable through expert curation and adaptable strategies. Importantly, the benefits far outweigh the costs: label transfer via UCE enables mostly consistent annotation across species without retraining, accelerating target qualification and enhancing translatability.

Beyond serving as a harmonized resource, our comprehensive landscape supports practical use cases in safety and pharmacology, exemplified here through analyses of tissue-specific expression, immunological features, and on-target, off-site effects of biologics, bringing together multiple disciplines. All annotations, metrics, and original labels are shared alongside this publication to promote transparency, reuse, and further development. Together, this work offers a powerful, extensible platform for integrative translational research, contributing to early drug development pipelines and the 3Rs, ultimately supporting safer therapeutic development.

## Data availability

All datasets used in this study are publicly available and their accession numbers described in Supplementary Table S1. Scripts used in this study are available in Gitlab (gitlab.com/genmab-public/macaca-fascicularis-single-cell-landscape). The single-cell object containing all studies, raw and normalized counts, as well as annotated cell labels (human-cynomolgus monkey, cynomolgus monkey-cynomolgus monkey and author’s original annotation) and UCE embeddings will be available on Zenodo (10.5281/zenodo.18172083, under review).

## Supplementary Figures

**Supplementary Figure S1:**
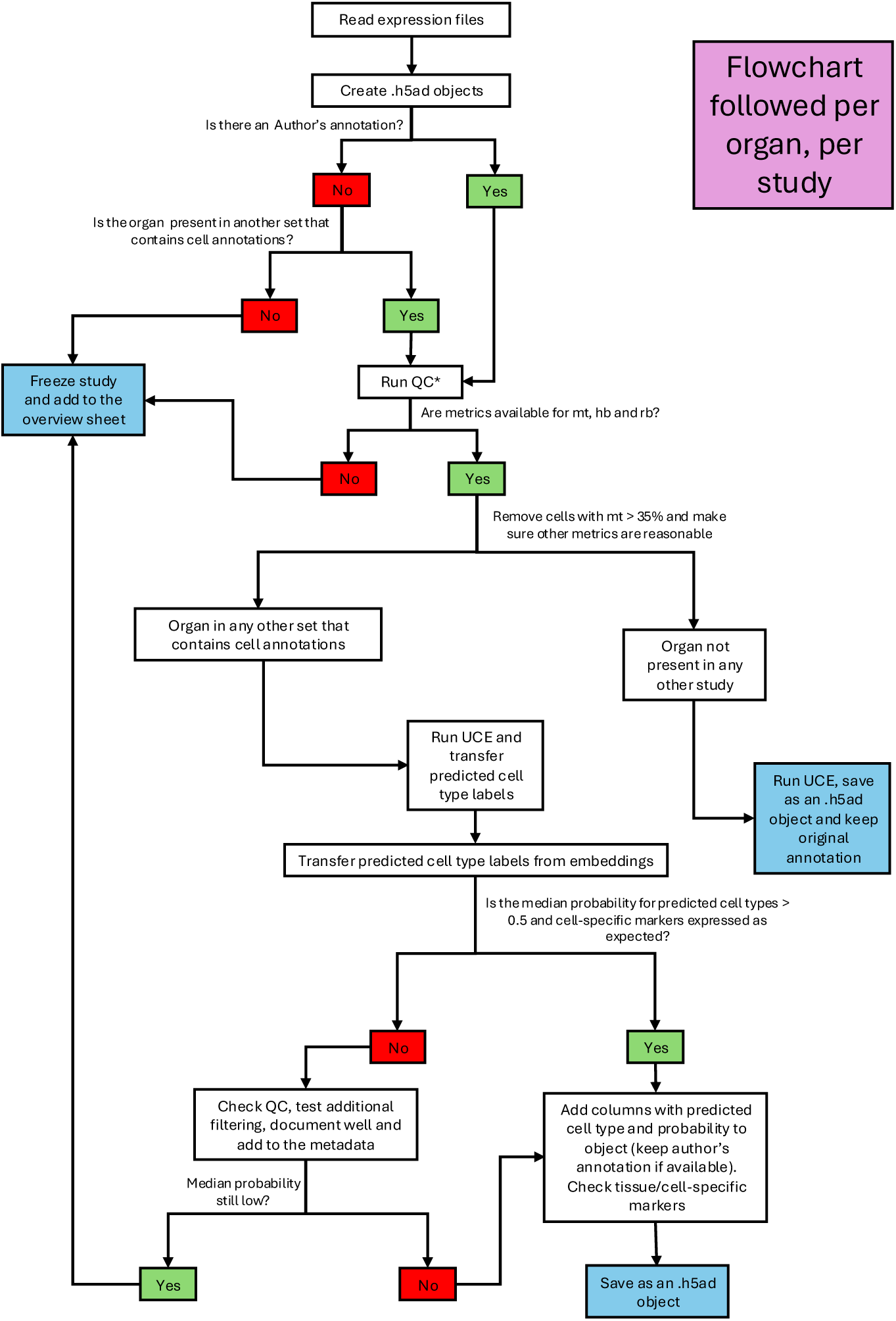
Data analysis workflow used for each single-cell/single-nuclei study. Flowchart used in the analysis of each study included. (*) Studies that did not have count tables available and were processed from .fastq files followed a stringent QC pipeline as described in the methods section of the manuscript.

**Supplementary Figure S2.**
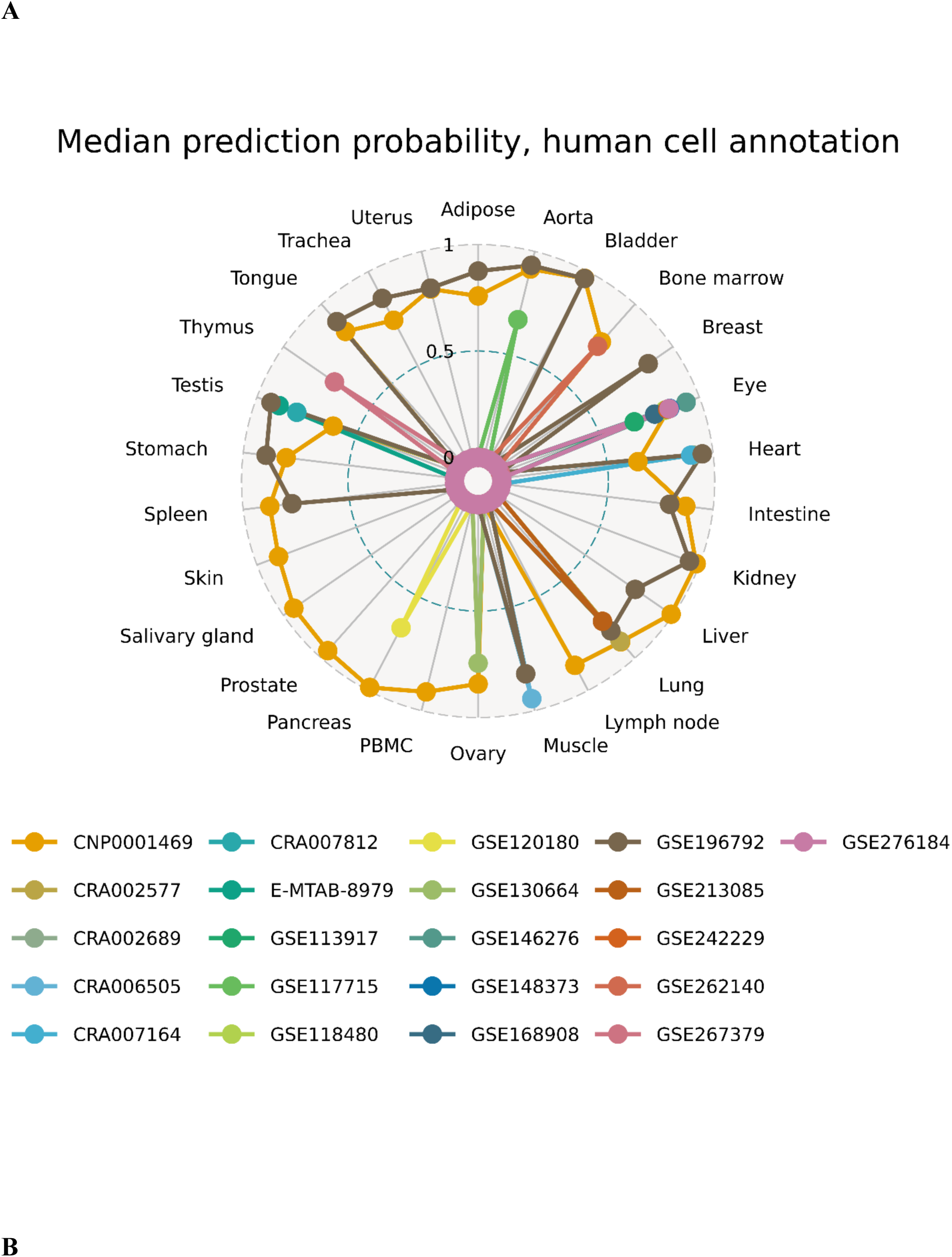
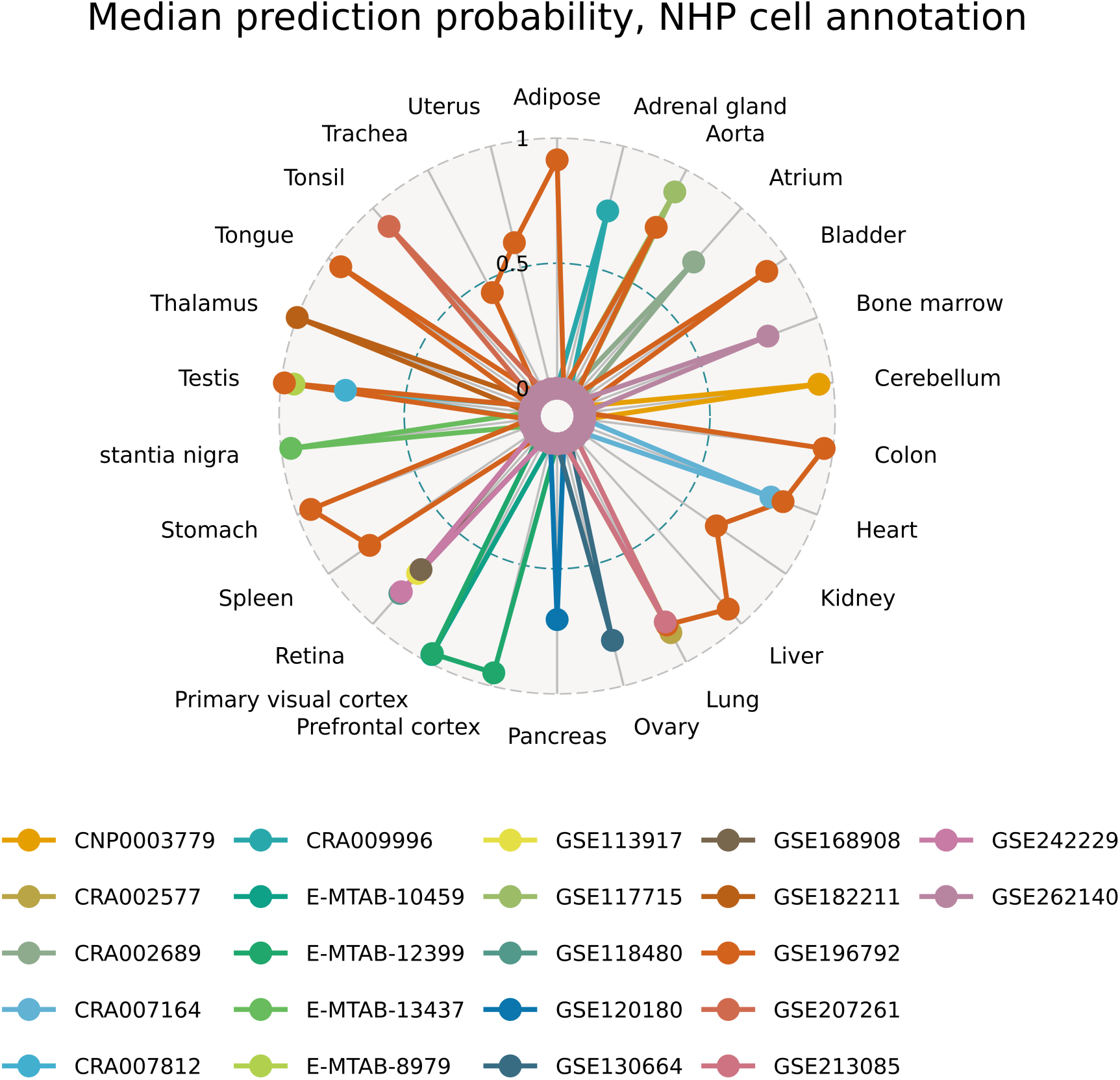
Median posterior probabilities after applying KNN to UCE embeddings. (A) Radar plots with median posterior probabilities per study across all organs using Tabula V2 as the reference. (B) Radar plots with median posterior probabilities per study across all organs using another non-human primate (NHP) study containing cell annotations as the reference (details in Supplementary Tables S1).

**Supplementary Figure S3.**
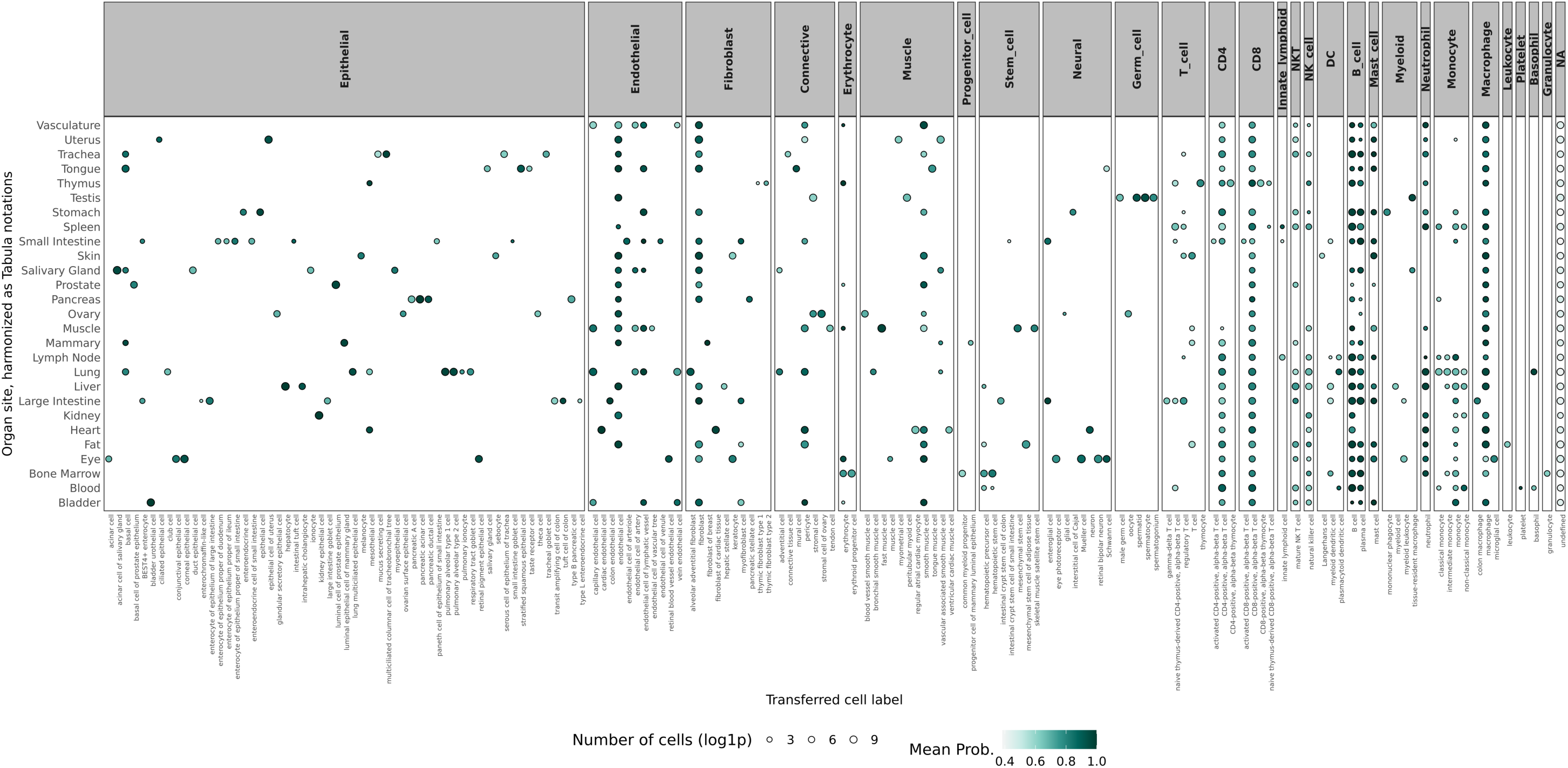
Median KNN posterior probabilities across cell types, organs and lineages in the non-human primate (NHP) *M. fascicularis* single cell landscape. Median posterior probabilities were calculated per cell type, across all studies included in each organ. Organs were harmonized to follow the same annotation as that present in Tabula V2, as presented in the column “tissue_in_publication”). Cell labels are based on KNN posterior prediction probabilities > 0.5, and those that did not surpass this threshold were labelled “undefined”.

**Supplementary Figure S4.**
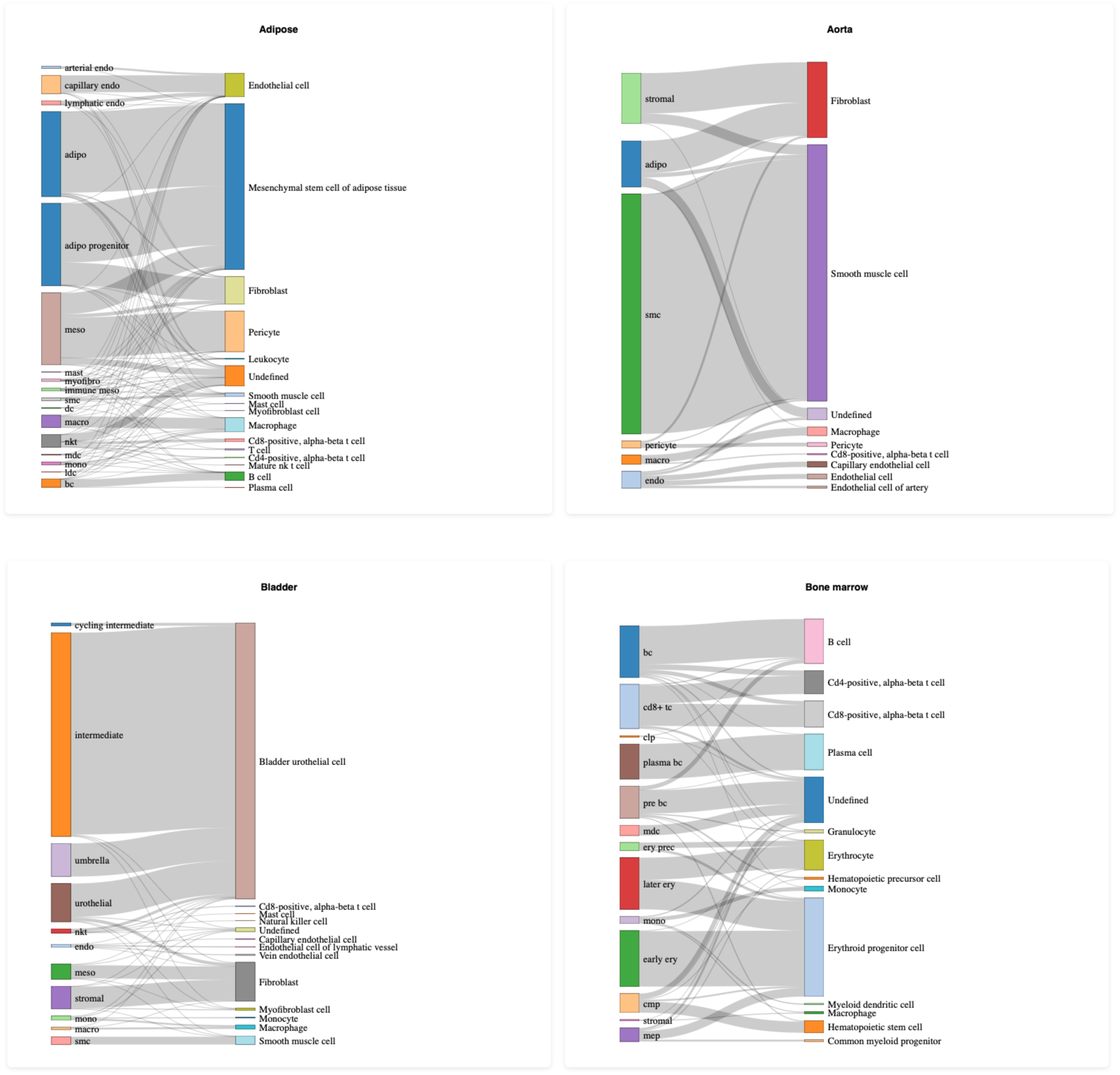
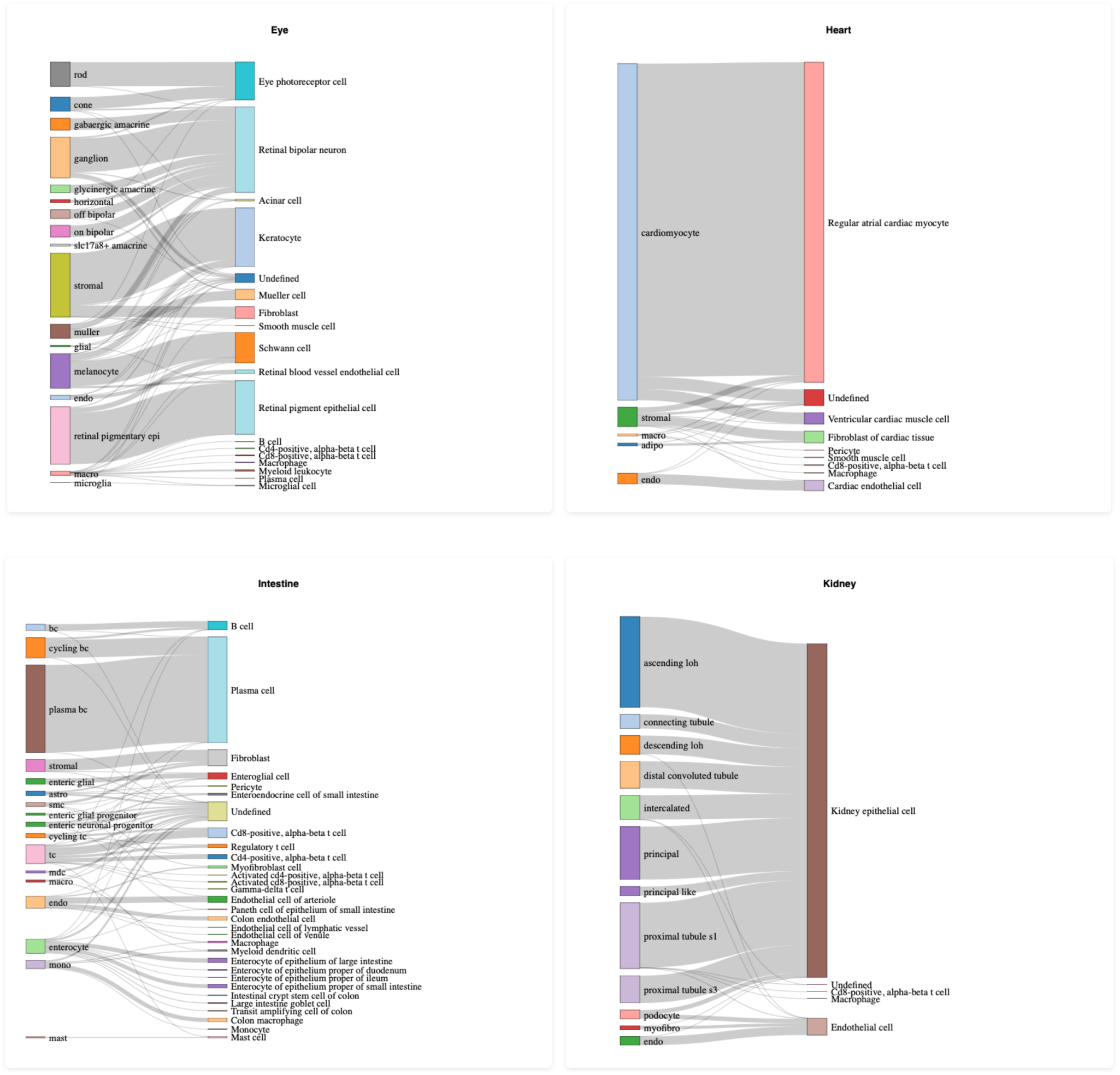
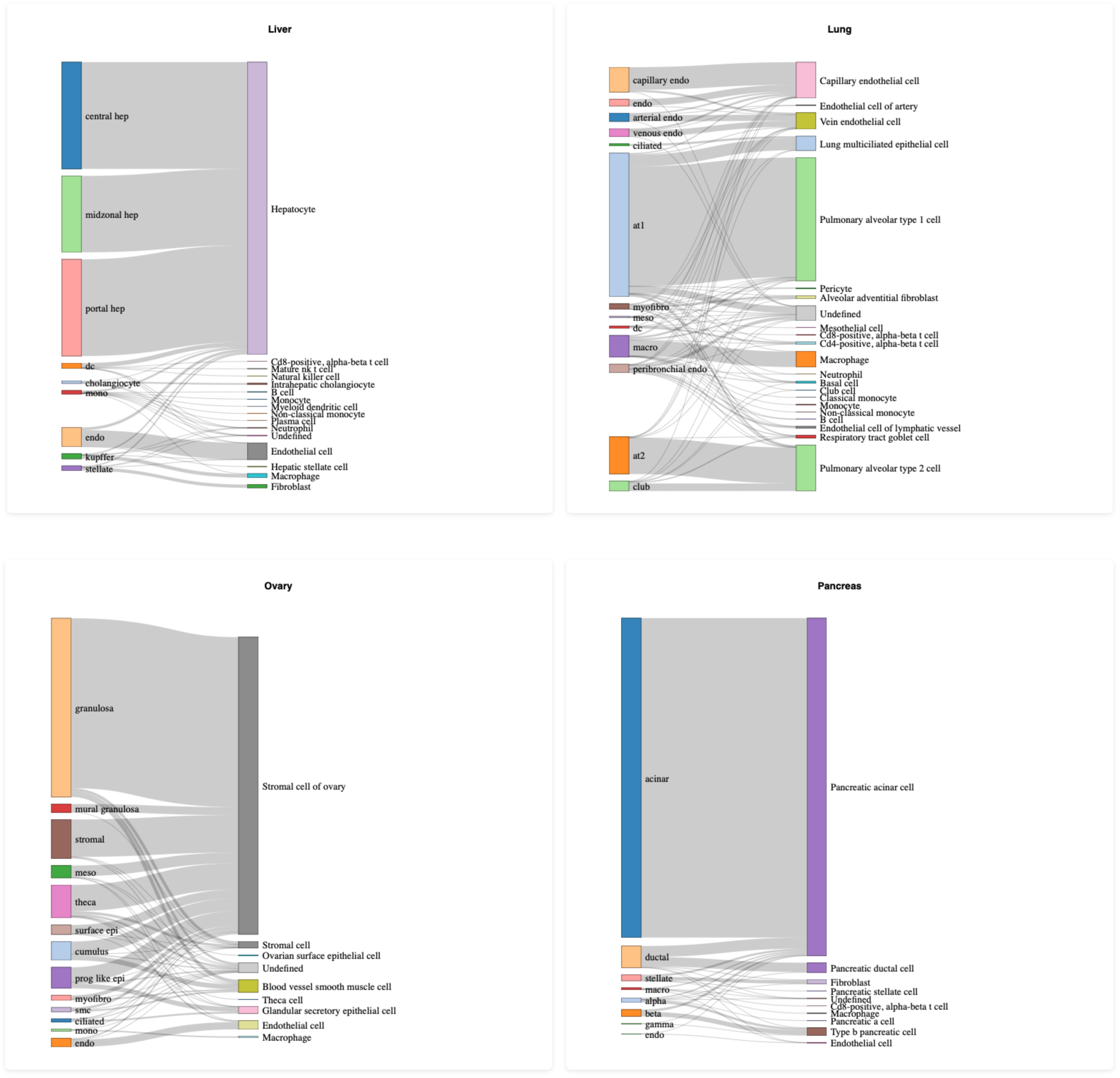
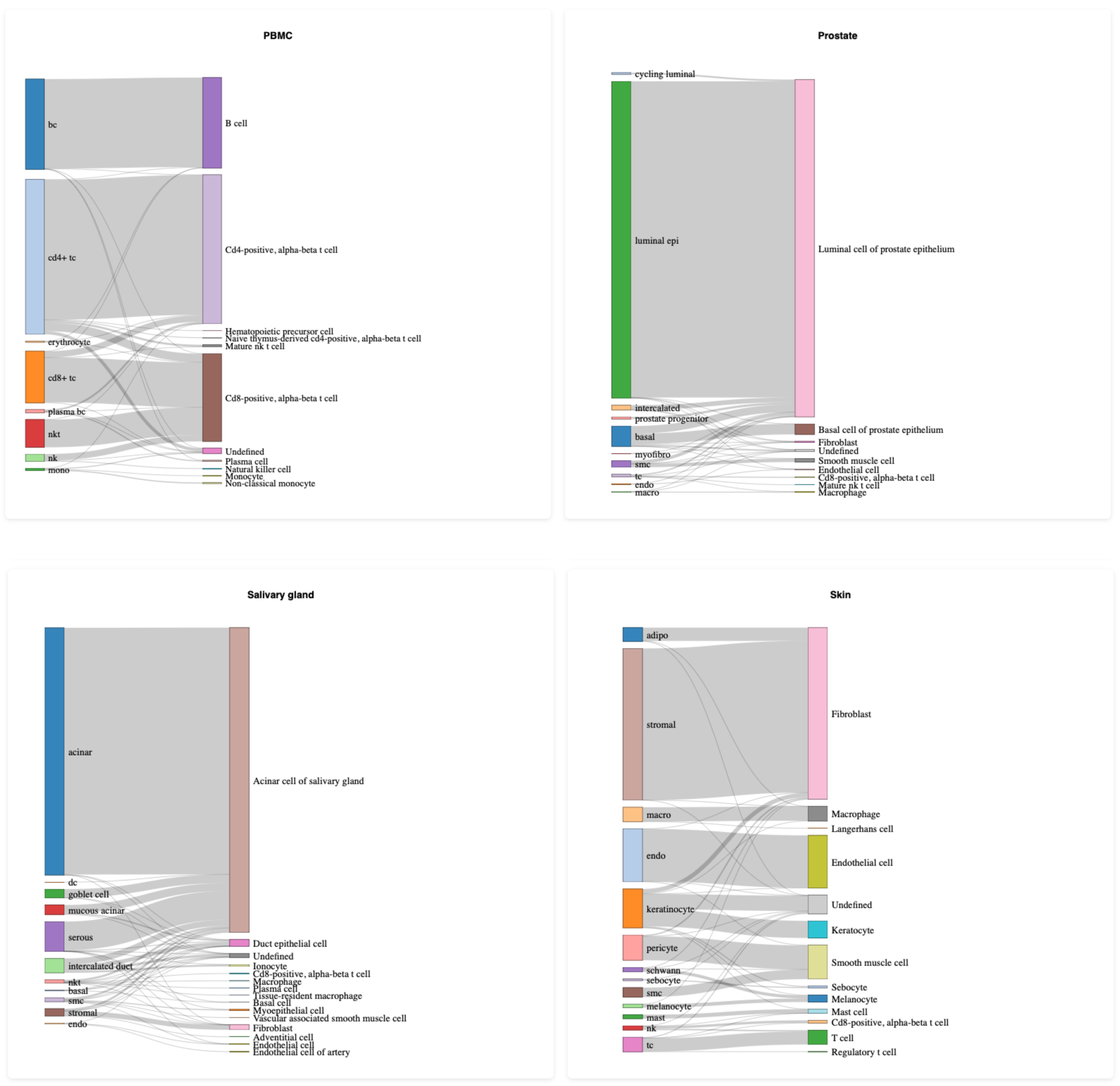
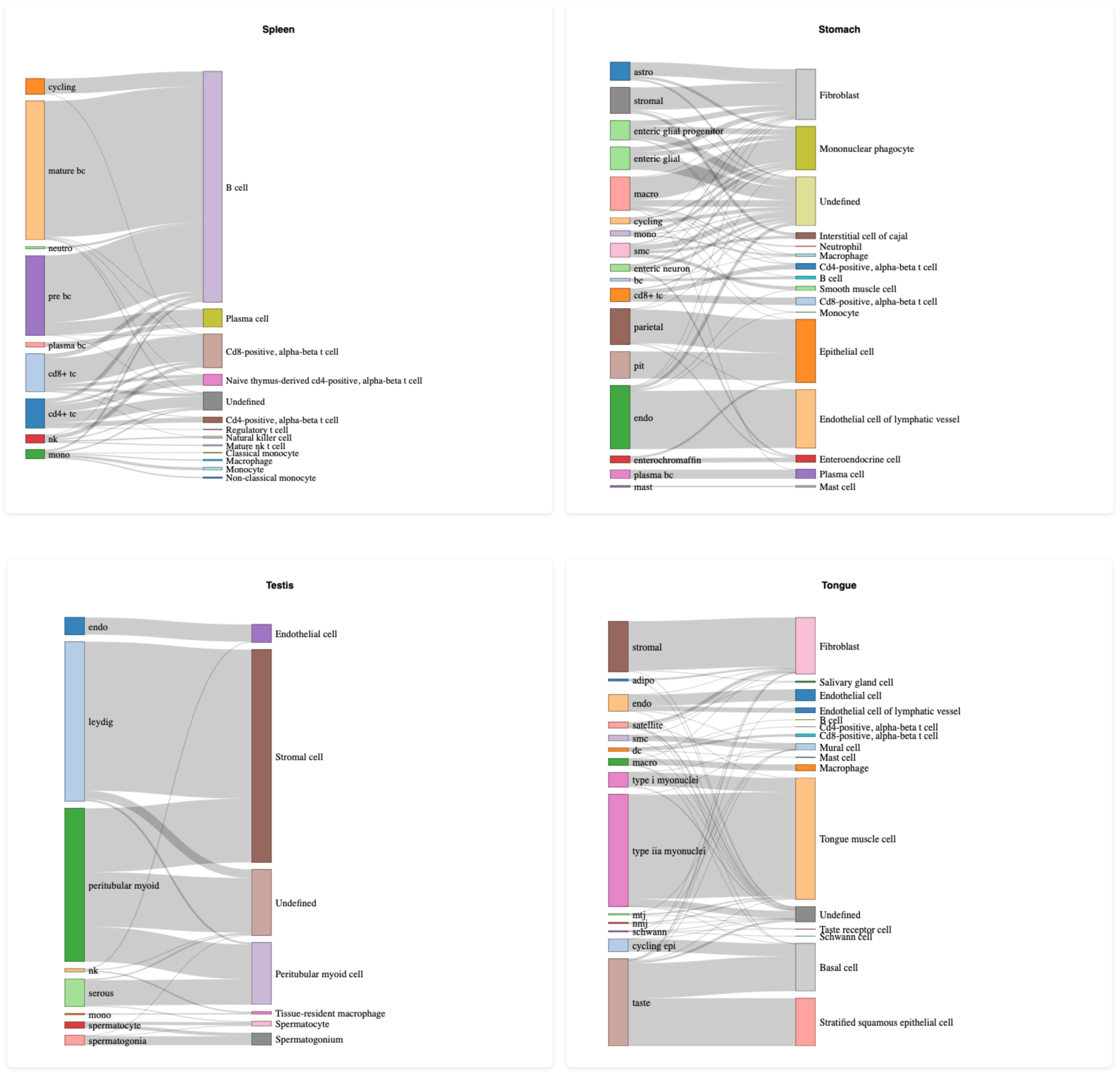
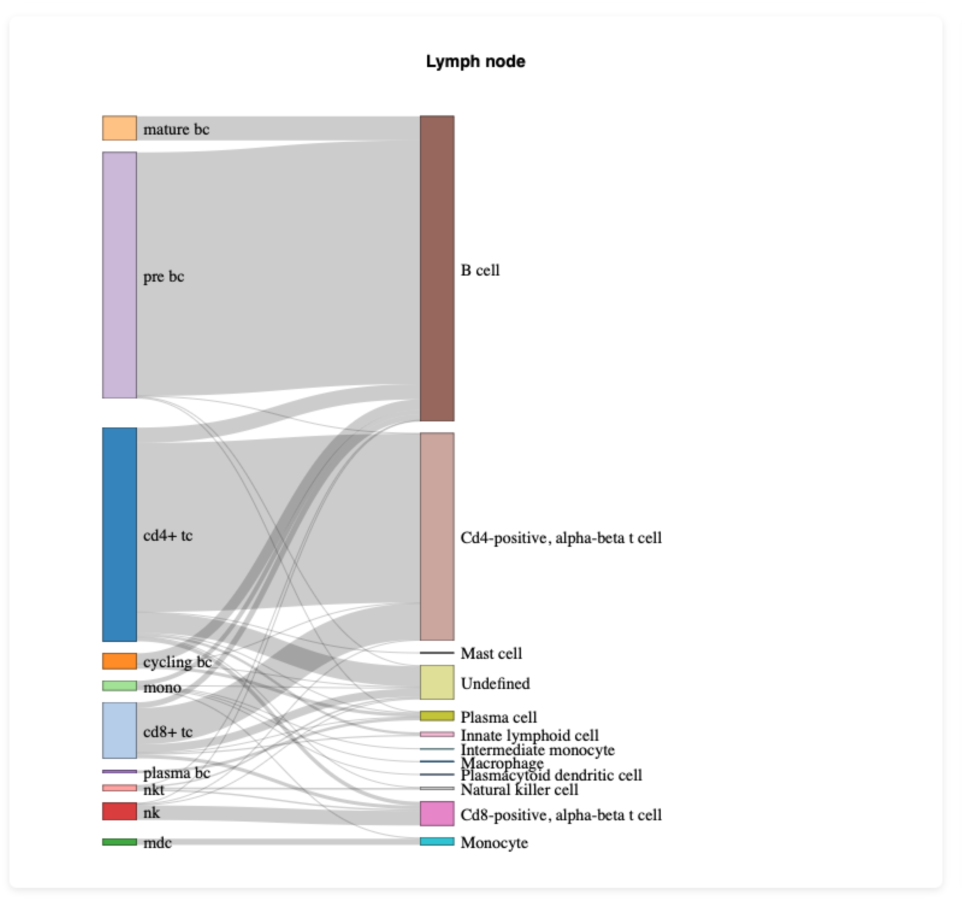
Sankey plots comparing original annotations from the Han et al., 2022 study (NHP, left) and cell labels predicted using Tabula V2 (human, right) as reference.

**Supplementary Figure S5.**
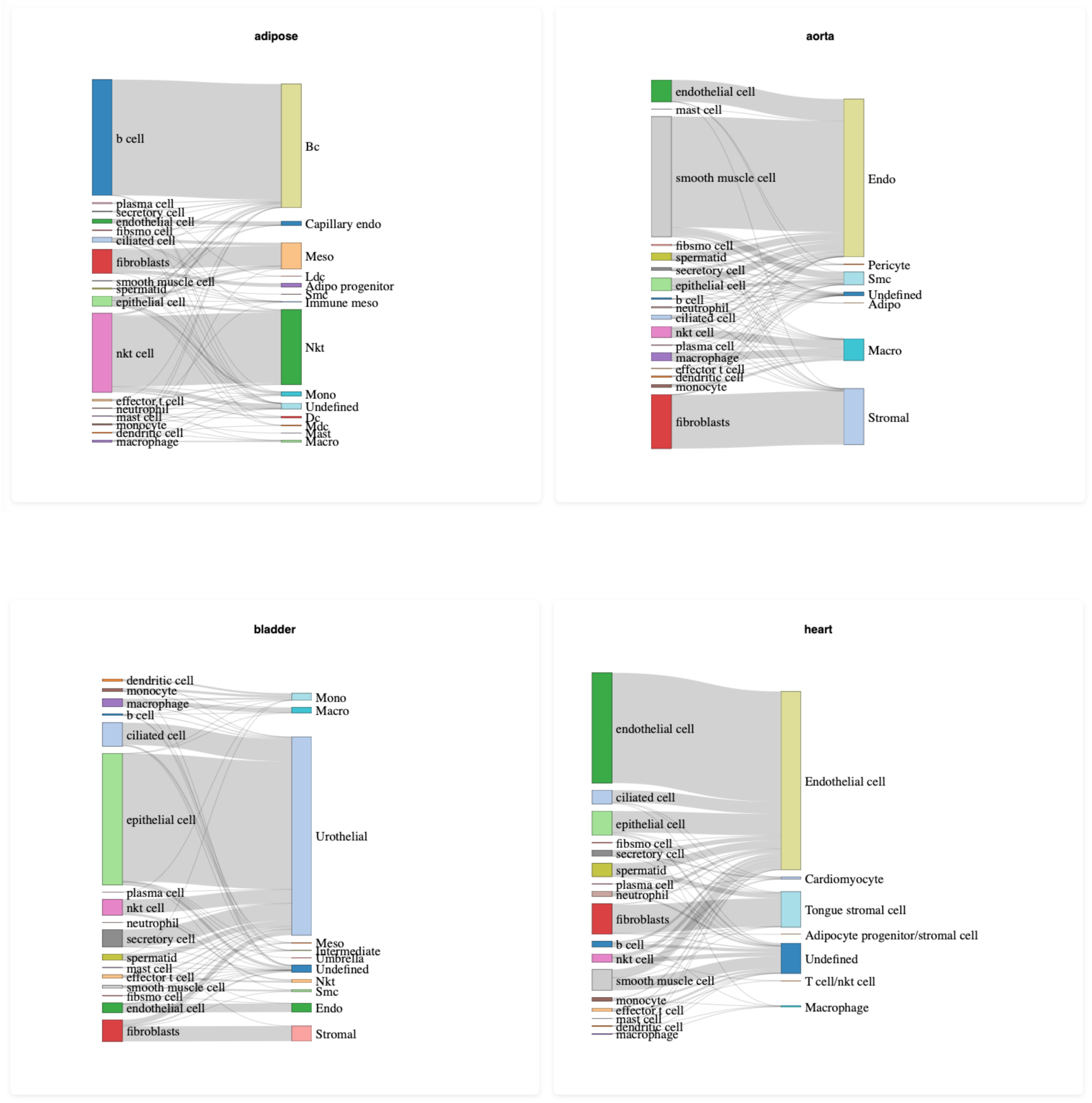
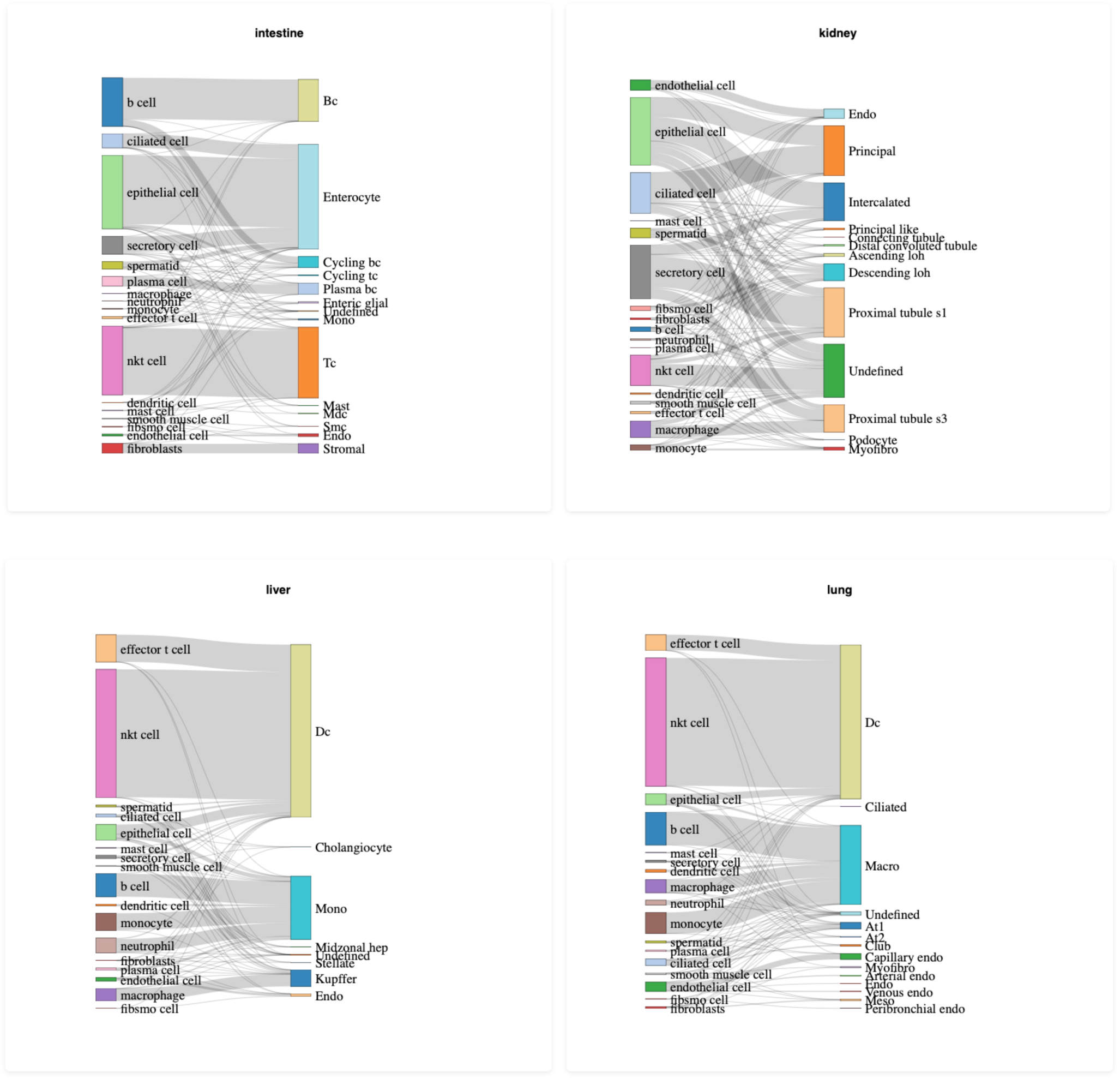
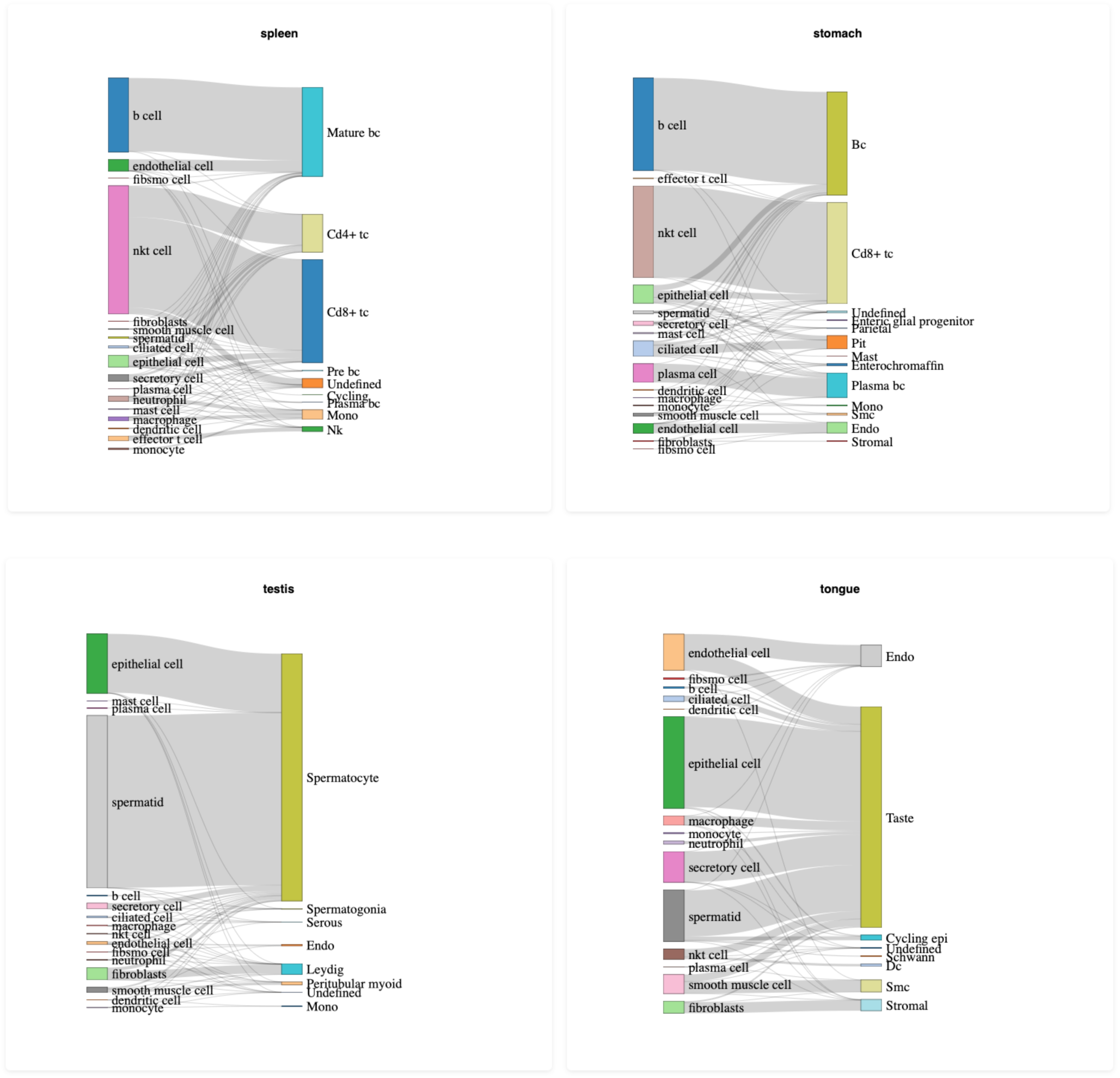
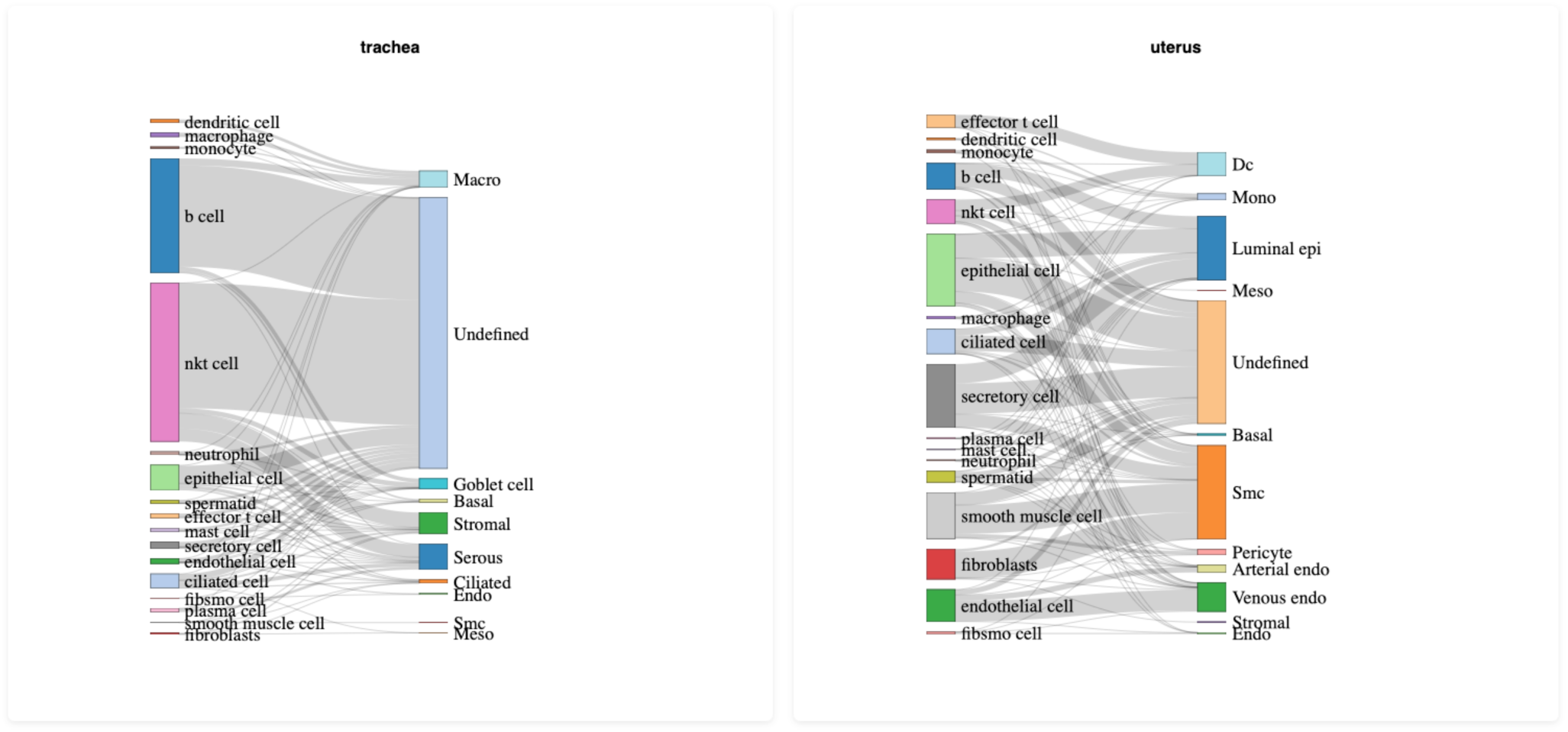
Sankey plots comparing original annotations from Qu et al. (2022) (NHP, left) and cell labels predicted using Han et al. (2021) (NHP, right) as reference.

**Supplementary Figure S6.**
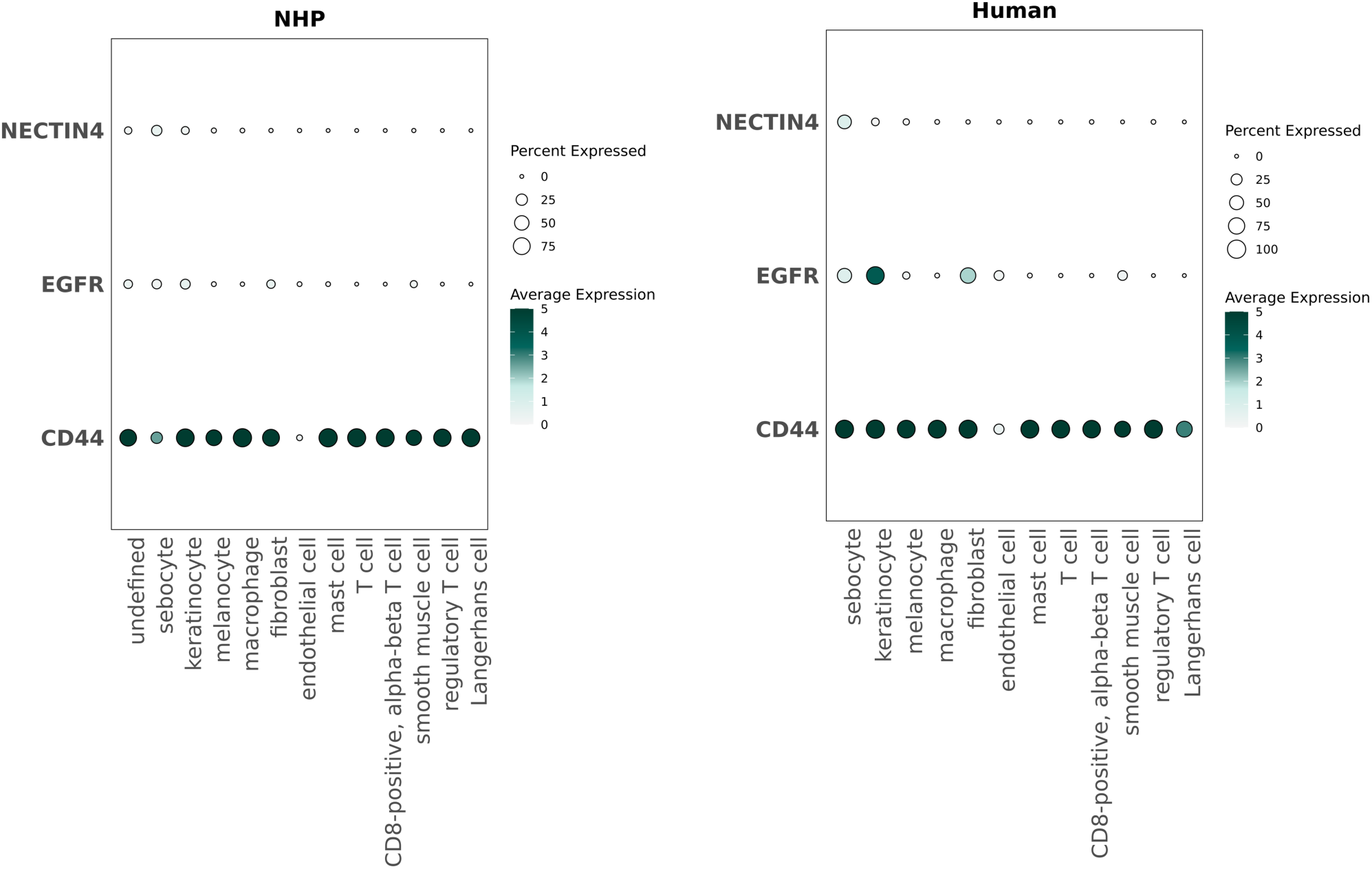
Dot plot depicting expression of CD44, EGFR and NECTIN4 in skin cell populations in non-human primate (NHP) and human. Cell labels were transferred from Tabula V2 following UCE after applying KNN to UCE embeddings. Cells labelled as “undefined” are those with posterior probability values of 0.5 or less.

**Supplementary Figure S7.**
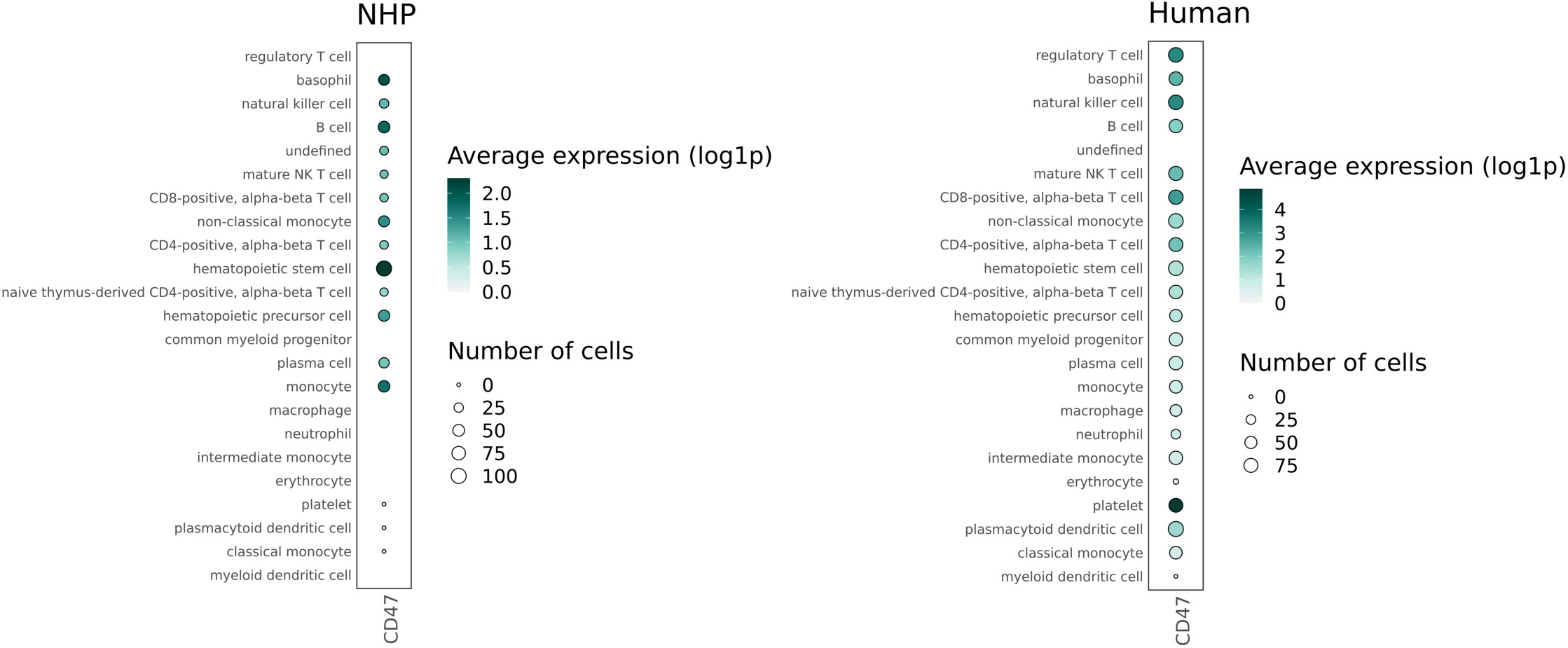
Dot plot depicting CD47 expression in blood samples from non-human primates (NHP) (PBMC, left) and human (blood fractions, right). Cell labels in NHPs were transferred from Tabula V2 following UCE after applying KNN to UCE embeddings. Cells labelled as “undefined” are those with posterior probability values of 0.5 or less.

